## Supplementary information for "Ecology, more than antibiotics consumption, is the major predictor for the global distribution of aminoglycoside-modifying enzymes"

**Text S1: Description of interaction effects on the prevalence of ACBs in European data between 1997 and 2018**

Aminoglycoside consumption can have a positive significant effect on the probability to sample ACBs in clinical samples (for CHG 6.1), soil (for CHG 6.1), farms samples (for CHGs 6.1 and 8), and domestic animals (for CHG 8), but also a negative significant effect in clinical samples (for CHG 17.1). Human exchanges also have ecology-specific impacts for 3 CHGs only. For CHG 6.1, migration has a positive effect on the probability to sample ACBs in clinical, farms, human, and soil samples but a negative effect in freshwater samples, whereas trade has opposite effects in all these biomes. For CHG 8, migration has a negative effect in soil samples, and trade has a negative effect in farms samples. For CHG 27, migration has a negative effect in clinical, farms, and human samples, while trade has a positive effect in farms.

**Text S2: Keywords used to filter sequences based on InterProScan annotations**

Amino acid sequences that were screened in genomes with HMMER AME profiles, were submitted to InterProScan in order to infer their putative functions. These sequences were then filtered based on keywords among their inferred functions. On the one hand, sequences that may have an enzymatic function similar to the ones of AMEs were screened with the following keywords: “acetyltransferase”, “adenylyltransferase”, “adenyltransferase”, “phosphotransferase”, “phosphoryltransferase”, and “nucleotidyltransferase”. On the other hand, sequences whose functions might be involved in aminoglycoside resistance were screened with the following keywords: “aminoglycoside”, “aminoside”, “mycin”, “micin”, and “amikacin”.

**Text S3: Keywords used to classify BioSamples into ecological biomes**

Among the BioSamples metadata columns, a number of columns were isolated to search for information enabling to characterize a sampling biome for each genome. On the one hand, were selected the column whose names contained one of the following keywords: “biome”, “habitat”, “env”, “host”. On the other hand, other columns were selected if they contained the keyword “source” or “isolat”, but none of the following keywords: “date”, “year”, “by”, “id”, “preservation”, “number”, “energy”, “carbon”, “method”, “comment”, “note”, “resource”, “history”, “annot”, or “lab”. Samples were then classified into biomes based on the contents of these columns.

Samples were classified as from clinical origin:

- if one of these columns (or columns regarding the sampling location) contained the keyword “hospit”, “clinic”, or “medical”;
- if one of these columns contained the keyword “human” or “homo” and at least one of the following keywords: “disease”, “blood”, “oral”, “feces”, “infection”, “arthr”, “hemo”, “bronch”, “pulmo”, “respi”, “urine”, “failure”, “pus”, “throat”, “thora”, “semen”, “wound”, “skin”, “gut”, “intestin”, “septicemia”, “sputum”, “fibrosis”, “swab”, “fluid”, “itis”, “bile”, “pneumonia”, “sputamentum”, “sick”, “gastr”, “aspirate”, “fecal”, “sudate”, “groin”, “emia”, “faecal”, “nares”, “osis”, “excreted”, “sepsis”, “patient”, “vagin”, “rect”, “surg”, “drainage”, “trach”, “lung”, “nasal”, “tissue”, “cornea”, “nosocomial”, “ICU”, “fever”, “bronc”, “absc”, “phary”;
- if the host was not specified but one of these columns contained any of the keywords: “melioidosis”, “tract infection”, “burn”, “urine”, “feces”, “blood”, “stool”, “sputum”

Samples were classified as originating from human habitats if they were not already assigned to clinical samples, and if their columns contained any of the following: “habitat”, “rural”, “bath”, “potable”, “tap water”, “toilet”, “spacecraft”, “food”.

Samples were classified as originating from domestic animals if they were not already assigned to clinical or human habitat samples, and if their columns contained any of the following: “canis”, “canine”, “feline”, “dog” (but not “hot-dog”) or “prairie dog”), “felis”, “rattus”, “cavia”, “cricet”, “chinchilla”, “mustela”, “mus”, “serinus”, “tortoise”, “parrot”, “carassius”, “mouse”.

Samples were classified as originating from farms if they were not already assigned to any of the previous origins, and if their columns contained any of the following: “bos”, “livestock”, “ovis”, “ovine”, “sheep”, “capra”, “meat”, “anas”, “goose”, “duck”, “cow”, “ovine”, “fish farm”, “slaughterhouse”, “poultry”, “gallus”, “pig”, “pork”, “beef”, “meleagris”, “oyster” (if also containing “cult”), “mellifera”, “milk”, “hen”, “chicken”, “lamb”, “egg”, “cattle”, “turkey”, “swine”, “porcine”, “goat”, “equine”, “horse”, “rabbit”, “calf”, “sus scrofa”, “equus”, “oryctolagus”, “manure”, “dairy”, “chease”, “sus” (if also containing “disease”), “farm”.

Samples were classified as originating from agrosystems if they were not already assigned to any of the previous origins, and if their columns contained any of the following: “wheat”, “salad”, “triticum”, “maize”, “zea mays”, “rice”, “oryza”, “field”, “plantation”, “olive”, “coffee”, “coffea”, “bean”, “banana”, “tomato”, “potato”, “solanum”, “musa”, “avena”, “oats”, “soy”, “brassica”, “hordeum”, “barley”, “rye”, “sorghum”, “millet”, “phaseolus”, “saccharum, “quinoa”, “cicer”, “pisum”, “sesam”, “cassava”, “sugar”, “manihot”, “ipomo”, “coconut”, “cotton”, “oil palm”, “helianthus”, “berry”, “onion”, “grape”, “walnut”, “peanut”, “prunus”, “citrus”, “greenhouse”, “lentil”, “tea”, “camellia”, “compost”, “tobacco”, “nicotiana”, “lettuce”, “vegetable”, “carrot”, “cucumber”, “pepper”, “pea”, “cilantro”.

Samples were classified as originating from wild plants and animals if they were not already assigned to any of the previous origins, and if their columns contained any of the following: “tapir”, “panthera”, “forest”, “grass”, “prairie dog”, “snake”, “ursus”, “ursi”, “migratory”, “crow”, “pika”, “gull”, “bird”, “clover”, “grazing”, “monkey”, “elephant”, “corvus”, “zebra”, “lily”, “vulture”.

Samples were classified as originating from freshwater if they were not already assigned to any of the previous origins, and if their columns contained any of the following: “river”, “marsh”, “pond”, “lake”, “swamp”, “bog”, “glacier”, “freshwater”, “fresh water”, “aquifer”, “permafrost”, “ground water”, “wetland”, “spring”, “catfish”, “creek”, “mineral water”, “environmental (streams)”.

Samples were classified as originating from sea water if they were not already assigned to any of the previous origins, and if their columns (or columns regarding the sampling location) contained any of the following: “ocean”, “seawater”, “sea”, “sea water”, “hydrotherm”, “continental shelf”, “saltwater”, “salt water”, “algae”, “tidal”, “tuna”, “seafood”, “shrimp”. These samples were later discarded because their sampling location could not be assigned to countries.

Samples were classified as originating from waste and sludge if they were not already assigned to any of the previous origins, and if their columns contained any of the following: “wastewater”, “waste water”, “sewage”, “sludge”.

Samples were classified as originating from soil if they were not already assigned to any of the previous origins, and if their columns contained any of the following: “soil”, “terrestrial”, “sand”, “humus”.

Finally, among the remaining samples that were assigned to none of the previous origins, samples whose columns contained the keywords “homo sapiens” or “human” were considered as originating from humans.

**Table S3: Summary of the selected model for CHG 1.** φ correspond to the estimated time autoregression coefficient. ν and ρ are respectively the estimated smoothness and scaling parameters of the Matérn correlation function. *: $p<0.05$, **: $p<0.01$, ***: $p<0.001$.

φ = -0.205, ν = 0.196, ρ = 0.36

| **Variable** | **Estimate** | **Conditional standard error** | **t** | **p** |
|---|---|---|---|---|
| Intercept | -6.11 | 1.54 | -3.98 | $6.97 \times 10^{-5}$ *** |
| Domestic animals | -34.2 | $5.96 \times 10^{6}$ | $-5.73 \times 10^{-6}$ | 1 |
| Farms | -5.34 | 0.576 | -9.26 | 0 *** |
| Flora, fauna | -33.3 | $8.04 \times 10^{6}$ | $-4.14 \times 10^{-6}$ | 1 |
| Human samples | -32.8 | $2.06 \times 10^{6}$ | $-1.59 \times 10^{-5}$ | 1 |
| Human habitat | -29.7 | $1.51 \times 10^{7}$ | $-1.97 \times 10^{-6}$ | 1 |
| Freshwater | -1.93 | 1.27 | -1.52 | 0.129 |
| Clinical samples | -4.99 | 0.587 | -8.51 | 0 *** |
| Sludge, waste | -35.7 | $4.48 \times 10^{6}$ | $-7.96 \times 10^{-6}$ | 1 |
| Soil | -2.82 | 1.24 | -2.28 | $2.28 \times 10^{-2}$ * |
| Trade | $5.83 \times 10^{-2}$ | 0.308 | 0.189 | 0.85 |
| Migration | 0.829 | 0.589 | 1.41 | 0.159 |
| Aminoglycosides | 1.63 | 0.589 | 2.76 | $5.71 \times 10^{-3}$ ** |

**Table S4: Summary of the selected model for CHG 2.** φ correspond to the estimated time autoregression coefficient. ν and ρ are respectively the estimated smoothness and scaling parameters of the Matérn correlation function. *: $p<0.05$, **: $p<0.01$, ***: $p<0.001$.

$\varphi = 0.597$, $\nu = 0.192$, $\rho = 10.3$

| **Variable** | **Estimate** | **Conditional standard error** | **t** | **p** |
|---|---|---|---|---|
| Intercept | -9.34 | 1.12 | -8.33 | $1.11 \times 10^{-16}$ *** |
| Domestic animals | -28.4 | $6.23 \times 10^{6}$ | $-4.56 \times 10^{-6}$ | 1 |
| Farms | -28.5 | $1.61 \times 10^{6}$ | $-1.77 \times 10^{-5}$ | 1 |
| Flora, fauna | -29.6 | $8.95 \times 10^{6}$ | $-3.3 \times 10^{-6}$ | 1 |
| Human samples | -28.1 | $2.08 \times 10^{6}$ | $-1.35 \times 10^{-5}$ | 1 |
| Human habitat | -27.7 | $1.33 \times 10^{7}$ | $-2.08 \times 10^{-6}$ | 1 |
| Freshwater | -30 | $8.93 \times 10^{6}$ | $-3.36 \times 10^{-6}$ | 1 |
| Clinical samples | 1.93 | 0.756 | 2.55 | $1.08 \times 10^{-2}$ * |
| Sludge, waste | -27.9 | $4.4 \times 10^{6}$ | $-6.35 \times 10^{-6}$ | 1 |
| Soil | -28.9 | $6.17 \times 10^{6}$ | $-4.69 \times 10^{-6}$ | 1 |

**Table S5: Summary of the selected model for CHG 3.** φ correspond to the estimated time autoregression coefficient. ν and ρ are respectively the estimated smoothness and scaling parameters of the Matérn correlation function. *: $p<0.05$, **: $p<0.01$, ***: $p<0.001$.

$\varphi = 7.59 \times 10^{-8}$, $\nu = 16.7$, $\rho = 8.47$

| **Variable** | **Estimate** | **Conditional standard error** | **t** | **p** |
|---|---|---|---|---|
| *Intercept* | -13.7 | 1.52 | -9.02 | 0 *** |
| *Domestic animals* | -26.9 | $7.9 \times 10^{6}$ | $-3.4 \times 10^{-6}$ | 1 |
| *Farms* | -25.5 | $1.19 \times 10^{5}$ | $-2.14 \times 10^{-4}$ | 1 |
| *Flora, fauna* | -25.3 | $1.37 \times 10^{6}$ | $-1.86 \times 10^{-5}$ | 1 |
| *Human samples* | 3.52 | 1.19 | 2.96 | $3.05 \times 10^{-3}$ ** |
| *Human habitat* | -22.6 | $1.15 \times 10^{7}$ | $-1.97 \times 10^{-6}$ | 1 |
| *Freshwater* | 4.46 | 1.79 | 2.5 | $1.26 \times 10^{-2}$ * |
| *Clinical samples* | 6.45 | 1.07 | 6.05 | $1.41 \times 10^{-9}$ *** |
| *Sludge, waste* | -28 | $4.51 \times 10^{6}$ | $-6.2 \times 10^{-6}$ | 1 |
| *Soil* | 4.41 | 1.48 | 2.98 | $2.85 \times 10^{-3}$ ** |
| *Trade* | -0.952 | 0.307 | -3.1 | $1.91 \times 10^{-3}$ ** |
| *Migration* | 1.73 | 0.582 | 2.97 | $2.95 \times 10^{-3}$ ** |
| *Aminoglycosides* | 1.48 | 0.124 | 12 | 0 *** |

**Table S6: Summary of the selected model for CHG 4.** φ correspond to the estimated time autoregression coefficient. ν and ρ are respectively the estimated smoothness and scaling parameters of the Matérn correlation function. *: $p<0.05$, **: $p<0.01$, ***: $p<0.001$.

$\varphi = 3.56 \times 10^{-4}$, $\nu = 6.04$, $\rho = 1.33$

| **Variable** | **Estimate** | **Conditional standard error** | **t** | **p** |
|---|---|---|---|---|
| Intercept | -36 | $3.42 \times 10^{6}$ | $-1.05 \times 10^{-5}$ | 1 |
| Domestic animals | -2.92 | $6.77 \times 10^{6}$ | $-4.32 \times 10^{-7}$ | 1 |
| Farms | 29.7 | $3.42 \times 10^{6}$ | $8.68 \times 10^{-6}$ | 1 |
| Flora, fauna | 31.6 | $3.42 \times 10^{6}$ | $9.24 \times 10^{-6}$ | 1 |
| Human samples | 29.1 | $3.42 \times 10^{6}$ | $8.51 \times 10^{-6}$ | 1 |
| Human habitat | -2.03 | $1.12 \times 10^{7}$ | $-1.81 \times 10^{-7}$ | 1 |
| Freshwater | -3.41 | $9.15 \times 10^{6}$ | $-3.73 \times 10^{-7}$ | 1 |
| Clinical samples | 29.8 | $3.42 \times 10^{6}$ | $8.71 \times 10^{-6}$ | 1 |
| Sludge, waste | -2 | $5.54 \times 10^{6}$ | $-3.61 \times 10^{-7}$ | 1 |
| Soil | -2.63 | $6.6 \times 10^{6}$ | $-3.99 \times 10^{-7}$ | 1 |
| Trade | 1.33 | 0.225 | 5.9 | $3.72 \times 10^{-9}$ *** |
| Migration | -1.38 | 0.339 | -4.06 | $4.81 \times 10^{-5}$ *** |
| Aminoglycosides | 0.568 | 0.214 | 2.65 | $8.03 \times 10^{-3}$ ** |

**Table S7: Summary of the selected model for CHG 5.1.** φ correspond to the estimated time autoregression coefficient. ν and ρ are respectively the estimated smoothness and scaling parameters of the Matérn correlation function. *: $p<0.05$, **: $p<0.01$, ***: $p<0.001$.

φ = 0.557, ν = 16.7, ρ = 3.6

| Variable | Estimate | Conditional standard error | t | p |
|---|---|---|---|---|
| Intercept | -6.3 | 1.72 | -3.66 | $2.47 \times 10^{-4}$ *** |
| Domestic animals | -65.2 | $1.66 \times 10^{6}$ | $-3.92 \times 10^{-5}$ | 1 |
| Farms | -2.36 | 1.95 | -1.21 | 0.225 |
| Flora, fauna | -3.09 | 6.53 | -0.472 | 0.637 |
| Human samples | 2.46 | 1.67 | 1.48 | 0.139 |
| Human habitat | $-7.11 \times 10^{2}$ | $1.29 \times 10^{6}$ | $-5.5 \times 10^{-4}$ | 1 |
| Freshwater | -30.8 | $8.85 \times 10^{6}$ | $-3.48 \times 10^{-6}$ | 1 |
| Clinical samples | 3.74 | 1.65 | 2.27 | $2.34 \times 10^{-2}$ * |
| Sludge, waste | 1.77 | 2.31 | 0.766 | 0.444 |
| Soil | -30.3 | $7.39 \times 10^{6}$ | $-4.1 \times 10^{-6}$ | 1 |
| Trade | 0.802 | 1.69 | 0.474 | 0.635 |
| Migration | -2.15 | 2.76 | -0.781 | 0.435 |
| Domestic animals × Trade | 14.9 | $5.95 \times 10^{5}$ | $2.51 \times 10^{-5}$ | 1 |
| Farms × Trade | 3 | 1.88 | 1.6 | 0.11 |
| Flora, fauna × Trade | -16.4 | 10.8 | -1.51 | 0.13 |
| Human samples × Trade | -0.366 | 1.7 | -0.215 | 0.83 |
| Human habitat × Trade | $8.32 \times 10^{2}$ | $1.5 \times 10^{6}$ | $5.54 \times 10^{-4}$ | 1 |
| Freshwater × Trade | -0.792 | $1.19 \times 10^{7}$ | $-6.64 \times 10^{-8}$ | 1 |
| Clinical samples × Trade | -0.807 | 1.69 | -0.479 | 0.632 |
| Sludge, waste × Trade | -1.31 | 1.84 | -0.712 | 0.477 |
| Soil × Trade | -1.1 | $8.88 \times 10^{6}$ | $-1.23 \times 10^{-7}$ | 1 |
| Domestic animals × Migration | 3.86 | $7.45 \times 10^{5}$ | $5.18 \times 10^{-6}$ | 1 |
| Farms × Migration | -1.98 | 3 | -0.662 | 0.508 |
| Flora, fauna × Migration | 3.28 | 3.13 | 1.05 | 0.295 |
| Human samples × Migration | 1.75 | 2.76 | 0.632 | 0.528 |
| Human habitat × Migration | $-1.17 \times 10^{3}$ | $2.11 \times 10^{6}$ | $-5.54 \times 10^{-4}$ | 1 |
| Freshwater × Migration | 2.1 | $1.13 \times 10^{7}$ | 1.86 × 10-7 | 1 |
| Clinical samples × Migration | 1.94 | 2.75 | 0.703 | 0.482 |
| Sludge, waste × Migration | 3.16 | 3.11 | 1.01 | 0.31 |
| Soil × Migration | 2.03 | $8.93 \times 10^{6}$ | $2.27 \times 10^{-7}$ | 1 |

**Table S8: Summary of the selected model for CHG 6.1.** φ correspond to the estimated time autoregression coefficient. ν and ρ are respectively the estimated smoothness and scaling parameters of the Matérn correlation function. *: $p<0.05$, **: $p<0.01$, ***: $p<0.001$.

$\varphi = 0.971$, $\nu = 0.21$, $\rho = 9.39 \times 10^{-2}$

| **Variable** | **Estimate** | **Conditional standard error** | **t** | **p** |
|---|---|---|---|---|
| Intercept | -5 | 1.82 | -2.75 | $5.93 \times 10^{-3}$ ** |
| Domestic animals | $-2.14 \times 10^{2}$ | $1.64 \times 10^{6}$ | $-1.31 \times 10^{-4}$ | 1 |
| Farms | 0.811 | 0.62 | 1.31 | 0.191 |
| Flora, fauna | -33 | $1.02 \times 10^{7}$ | $-3.25 \times 10^{-6}$ | 1 |
| Human samples | 1.35 | 0.636 | 2.13 | $3.34 \times 10^{-2}$ * |
| Human habitat | 2.55 | 0.999 | 2.55 | $1.07 \times 10^{-2}$ * |
| Freshwater | -3.14 | 2.09 | -1.5 | 0.132 |
| Clinical samples | -0.795 | 0.62 | -1.28 | 0.2 |
| Sludge, waste | -3.44 | 6.39 | -0.537 | 0.591 |
| Soil | 0.721 | 0.88 | 0.82 | 0.412 |
| Trade | 1.5 | 0.648 | 2.31 | $2.11 \times 10^{-2}$ * |
| Migration | -1.06 | 0.501 | -2.12 | $3.4 \times 10^{-2}$ * |
| Aminoglycosides | -2.55 | 0.901 | -2.83 | $4.67 \times 10^{-3}$ ** |
| Domestic animals × Trade | 59.7 | $7.05 \times 10^{5}$ | $8.48 \times 10^{-5}$ | 1 |
| Farms × Trade | -2.49 | 0.618 | -4.03 | $5.66 \times 10^{-5}$ *** |
| Flora, fauna × Trade | -0.179 | $1.33 \times 10^{7}$ | $-1.35 \times 10^{-8}$ | 1 |
| Human samples × Trade | -2.12 | 0.602 | -3.52 | $4.36 \times 10^{-4}$ *** |
| Human habitat × Trade | -0.994 | 1.09 | -0.916 | 0.36 |
| Freshwater × Trade | 3.51 | 1.68 | 2.09 | $3.64 \times 10^{-2}$ * |
| Clinical samples × Trade | -2.07 | 0.612 | -3.38 | $7.23 \times 10^{-4}$ *** |
| Sludge, waste × Trade | -7.3 | 7.41 | -0.985 | 0.324 |
| Soil × Trade | -3.01 | 0.829 | -3.62 | $2.89 \times 10^{-4}$ *** |
| Domestic animals × Migration | 51.9 | $1.18 \times 10^{6}$ | $4.41 \times 10^{-5}$ | 1 |
| Farms × Migration | 2.03 | 0.49 | 4.15 | $3.29 \times 10^{-5}$ *** |
| Flora, fauna × Migration | 0.553 | $9.46 \times 10^{6}$ | $5.84 \times 10^{-8}$ | 1 |
| Human samples × Migration | 2.02 | 0.518 | 3.91 | $9.16 \times 10^{-5}$ *** |
| Human habitat × Migration | 0.725 | 1.36 | 0.532 | 0.595 |
| Freshwater × Migration | -3.87 | 1.56 | -2.47 | $1.35 \times 10^{-2}$ * |
| Clinical samples × Migration | 1.61 | 0.507 | 3.17 | $1.51 \times 10^{-3}$ ** |
| Sludge, waste × Migration | 2.33 | 1.87 | 1.25 | 0.212 |
| Soil × Migration | 2.2 | 0.944 | 2.33 | $1.97 \times 10^{-2}$ * |
| Domestic animals × Aminoglycosides | $-1.43 \times 10^{2}$ | $2.2 \times 10^{6}$ | $-6.51 \times 10^{-5}$ | 1 |
| Farms × Aminoglycosides | 2.57 | 0.92 | 2.79 | $5.28 \times 10^{-3}$ ** |
| Flora, fauna × Aminoglycosides | 1.52 | $1.46 \times 10^{7}$ | $1.04 \times 10^{-7}$ | 1 |
| Human samples × Aminoglycosides | 1.63 | 0.943 | 1.73 | $8.38 \times 10^{-2}$ * |
| Human habitat × Aminoglycosides | 1.13 | 1.65 | 0.683 | 0.495 |
| Freshwater × Aminoglycosides | -3.19 | 2.49 | -1.28 | 0.2 |
| Clinical samples × Aminoglycosides | 2.42 | 0.917 | 2.64 | $8.23 \times 10^{-3}$ ** |
| Sludge, waste × Aminoglycosides | -0.529 | 4.7 | -0.113 | 0.91 |

| | | | | |
|---|---|---|---|---|
| Soil × Aminoglycosides | 4.3 | 1 | 4.29 | $1.78 \times 10^{-5}$ *** |

**Table S9: Summary of the selected model for CHG 7.** φ correspond to the estimated time autoregression coefficient. ν and ρ are respectively the estimated smoothness and scaling parameters of the Matérn correlation function. *: $p<0.05$, **: $p<0.01$, ***: $p<0.001$.

φ = -0.429, ν = 16.7, ρ = 1.45

| Variable | Estimate | Conditional standard error | t | p |
|---|---|---|---|---|
| Intercept | -5.57 | 0.764 | -7.28 | $3.26 \times 10^{-13}$ *** |
| Domestic animals | -16.8 | 30.6 | -0.548 | 0.583 |
| Farms | $1.01 \times 10^{-2}$ | 0.76 | $1.33 \times 10^{-2}$ | 0.989 |
| Flora, fauna | $-6.75 \times 10^{2}$ | $8 \times 10^{6}$ | $-8.44 \times 10^{-5}$ | 1 |
| Human samples | 0.76 | 0.782 | 0.971 | 0.332 |
| Human habitat | $-3.86 \times 10^{2}$ | $3.6 \times 10^{6}$ | $-1.07 \times 10^{-4}$ | 1 |
| Freshwater | $-2.19 \times 10^{2}$ | $8.67 \times 10^{6}$ | $-2.52 \times 10^{-5}$ | 1 |
| Clinical samples | -0.126 | 0.714 | -0.176 | 0.86 |
| Sludge, waste | $-2.14 \times 10^{2}$ | $1.19 \times 10^{7}$ | $-1.8 \times 10^{-5}$ | 1 |
| Soil | -4.54 | 6.36 | -0.713 | 0.476 |
| Trade | $9.54 \times 10^{-2}$ | 0.339 | 0.281 | 0.779 |
| Migration | 0.34 | 0.519 | 0.656 | 0.512 |
| Aminoglycosides | 0.679 | 0.381 | 1.78 | $7.47 \times 10^{-2}$ * |
| Domestic animals × Trade | 5.97 | 10 | 0.597 | 0.551 |
| Farms × Trade | 0.465 | 0.363 | 1.28 | 0.2 |
| Flora, fauna × Trade | $-3.71 \times 10^{2}$ | $9.92 \times 10^{6}$ | $-3.74 \times 10^{-5}$ | 1 |
| Human samples × Trade | 0.528 | 0.363 | 1.45 | 0.146 |
| Human habitat × Trade | 68.6 | $6.51 \times 10^{5}$ | $1.05 \times 10^{-4}$ | 1 |
| Freshwater × Trade | 4.6 | $7.92 \times 10^{6}$ | $5.81 \times 10^{-7}$ | 1 |
| Clinical samples × Trade | 0.384 | 0.349 | 1.1 | 0.271 |
| Sludge, waste × Trade | 5.08 | $8.1 \times 10^{6}$ | $6.28 \times 10^{-7}$ | 1 |
| Soil × Trade | 1.55 | 2.04 | 0.761 | 0.447 |
| Domestic animals × Migration | 3.65 | 7.53 | 0.484 | 0.628 |
| Farms × Migration | -0.68 | 0.576 | -1.18 | 0.238 |
| Flora, fauna × Migration | $-2.61 \times 10^{2}$ | $7.39 \times 10^{6}$ | $-3.53 \times 10^{-5}$ | 1 |
| Human samples × Migration | -0.237 | 0.548 | -0.433 | 0.665 |
| Human habitat × Migration | $1.79 \times 10^{2}$ | $1.67 \times 10^{6}$ | $1.07 \times 10^{-4}$ | 1 |
| Freshwater × Migration | 3.51 | $9.19 \times 10^{6}$ | $3.82 \times 10^{-7}$ | 1 |
| Clinical samples × Migration | 0.558 | 0.51 | 1.1 | 0.273 |
| Sludge, waste × Migration | 2.89 | $1.2 \times 10^{7}$ | $2.41 \times 10^{-7}$ | 1 |
| Soil × Migration | $-3.87 \times 10^{-2}$ | 1.97 | $-1.96 \times 10^{-2}$ | 0.984 |
| Domestic animals × Aminoglycosides | 6.77 | 10.6 | 0.639 | 0.523 |
| Farms × Aminoglycosides | 0.176 | 0.497 | 0.355 | 0.723 |
| Flora, fauna × Aminoglycosides | 22.8 | $8.37 \times 10^{6}$ | $2.72 \times 10^{-6}$ | 1 |
| Human samples × Aminoglycosides | 0.134 | 0.426 | 0.314 | 0.754 |
| Human habitat × Aminoglycosides | $-2.8 \times 10^{2}$ | $2.68 \times 10^{6}$ | $-1.04 \times 10^{-4}$ | 1 |
| Freshwater × Aminoglycosides | 3.51 | $9.83 \times 10^{6}$ | $3.57 \times 10^{-7}$ | 1 |
| Clinical samples × Aminoglycosides | -2.15 | 0.524 | -4.1 | $4.11 \times 10^{-5}$ *** |
| Sludge, waste × Aminoglycosides | 3.47 | $1.88 \times 10^{7}$ | $1.85 \times 10^{-7}$ | 1 |
| Soil × Aminoglycosides | 1.26 | 2.2 | 0.574 | 0.566 |

**Table S10: Summary of the selected model for CHG 8.** φ correspond to the estimated time autoregression coefficient. ν and ρ are respectively the estimated smoothness and scaling parameters of the Matérn correlation function. *: p<0.05, **: p<0.01, ***: p<0.001.

$\varphi = 4.44 \times 10^{-2}$, $\nu = 0.701$, $\rho = 0.205$

| Variable | Estimate | Conditional standard error | t | p |
|---|---|---|---|---|
| Intercept | -4.95 | 0.639 | -7.74 | $1.01 \times 10^{-14}$ *** |
| Domestic animals | -13.1 | 7.71 | -1.69 | $9.04 \times 10^{-2}$ * |
| Farms | -1.41 | 0.645 | -2.19 | $2.87 \times 10^{-2}$ * |
| Flora, fauna | -3.97 | 6.36 | -0.624 | 0.532 |
| Human samples | 0.331 | 0.55 | 0.601 | 0.548 |
| Human habitat | -2.18 | 2.56 | -0.853 | 0.394 |
| Freshwater | -20.6 | $3.62 \times 10^{4}$ | $-5.68 \times 10^{-4}$ | 1 |
| Clinical samples | -0.764 | 0.516 | -1.48 | 0.138 |
| Sludge, waste | -2.86 | 4.7 | -0.608 | 0.543 |
| Soil | $6.16 \times 10^{-2}$ | 0.935 | $6.59 \times 10^{-2}$ | 0.947 |
| Trade | 1.42 | 0.638 | 2.23 | $2.57 \times 10^{-2}$ * |
| Migration | -0.434 | 0.801 | -0.542 | 0.588 |
| Aminoglycosides | -0.695 | 0.929 | -0.747 | 0.455 |
| Domestic animals × Trade | 6.73 | 3.94 | 1.71 | $8.78 \times 10^{-2}$ * |
| Farms × Trade | 0.151 | 0.697 | 0.216 | 0.829 |
| Flora, fauna × Trade | 1.84 | 4.92 | 0.375 | 0.708 |
| Human samples × Trade | -0.93 | 0.655 | -1.42 | 0.156 |
| Human habitat × Trade | 6.31 | 3.94 | 1.6 | 0.109 |
| Freshwater × Trade | -1.38 | $3.75 \times 10^{4}$ | $-3.67 \times 10^{-5}$ | 1 |
| Clinical samples × Trade | -0.883 | 0.641 | -1.38 | 0.168 |
| Sludge, waste × Trade | 2.59 | 13.2 | 0.196 | 0.844 |
| Soil × Trade | -4.7 | 1.9 | -2.47 | $1.34 \times 10^{-2}$ * |
| Domestic animals × Migration | -16.4 | 9.02 | -1.82 | $6.84 \times 10^{-2}$ * |
| Farms × Migration | -1.99 | 1 | -1.99 | $4.68 \times 10^{-2}$ * |
| Flora, fauna × Migration | -4.68 | 7.69 | -0.609 | 0.543 |
| Human samples × Migration | 0.248 | 0.834 | 0.298 | 0.766 |
| Human habitat × Migration | -9.79 | 5.03 | -1.95 | $5.17 \times 10^{-2}$ * |
| Freshwater × Migration | 0.484 | $5.19 \times 10^{4}$ | $9.33 \times 10^{-6}$ | 1 |
| Clinical samples × Migration | 0.99 | 0.805 | 1.23 | 0.219 |
| Sludge, waste × Migration | $3.63 \times 10^{-2}$ | 4.21 | $8.62 \times 10^{-3}$ | 0.993 |
| Soil × Migration | 1.47 | 0.884 | 1.67 | $9.57 \times 10^{-2}$ * |
| Domestic animals × Aminoglycosides | 6.28 | 3.18 | 1.98 | $4.82 \times 10^{-2}$ * |
| Farms × Aminoglycosides | 2.06 | 0.954 | 2.15 | $3.12 \times 10^{-2}$ * |
| Flora, fauna × Aminoglycosides | -4.24 | 7.09 | -0.597 | 0.55 |
| Human samples × Aminoglycosides | 1.37 | 0.935 | 1.46 | 0.144 |
| Human habitat × Aminoglycosides | -0.23 | 1.62 | -0.142 | 0.887 |
| Freshwater × Aminoglycosides | 0.649 | $3.07 \times 10^{4}$ | $2.12 \times 10^{-5}$ | 1 |
| Clinical samples × Aminoglycosides | -0.729 | 0.963 | -0.757 | 0.449 |
| Sludge, waste × Aminoglycosides | -3.34 | 17.6 | -0.19 | 0.849 |
| Soil × Aminoglycosides | 0.481 | 1.06 | 0.452 | 0.651 |

**Table S11: Summary of the selected model for CHG 11.** φ correspond to the estimated time autoregression coefficient. ν and ρ are respectively the estimated smoothness and scaling parameters of the Matérn correlation function. *: p<0.05, **: p<0.01, ***: p<0.001.

$\varphi = 0.368$, $\nu = 16.7$, $\rho = 1.49$

| **Variable** | **Estimate** | **Conditional standard error** | **t** | **p** |
|---|---|---|---|---|
| Intercept | -35.7 | $3.84 \times 10^{6}$ | $-9.28 \times 10^{-6}$ | 1 |
| Domestic animals | -48.9 | $6.94 \times 10^{6}$ | $-7.05 \times 10^{-6}$ | 1 |
| Farms | 29.5 | $3.84 \times 10^{6}$ | $7.68 \times 10^{-6}$ | 1 |
| Flora, fauna | -50.1 | $8.65 \times 10^{6}$ | $-5.79 \times 10^{-6}$ | 1 |
| Human samples | 30.9 | $3.84 \times 10^{6}$ | $8.05 \times 10^{-6}$ | 1 |
| Human habitat | -49.1 | $1.1 \times 10^{7}$ | $-4.44 \times 10^{-6}$ | 1 |
| Freshwater | -52.4 | $9.23 \times 10^{6}$ | $-5.68 \times 10^{-6}$ | 1 |
| Clinical samples | 29.7 | $3.84 \times 10^{6}$ | $7.73 \times 10^{-6}$ | 1 |
| Sludge, waste | -48.2 | $5.81 \times 10^{6}$ | $-8.29 \times 10^{-6}$ | 1 |
| Soil | -49.9 | $6.78 \times 10^{6}$ | $-7.35 \times 10^{-6}$ | 1 |
| Trade | 0.634 | 0.189 | 3.35 | $8 \times 10^{-4}$ *** |
| Migration | -0.342 | 0.329 | -1.04 | 0.299 |

**Table S12: Summary of the selected model for CHG 13.** φ correspond to the estimated time autoregression coefficient. ν and ρ are respectively the estimated smoothness and scaling parameters of the Matérn correlation function. *: $p<0.05$, **: $p<0.01$, ***: $p<0.001$.

φ = 0.874, ν = 16.7, ρ = 2.75

| Variable | Estimate | Conditional standard error | t | p |
|---|---|---|---|---|
| Intercept | -40.6 | $2.78 \times 10^{6}$ | $-1.46 \times 10^{-5}$ | 1 |
| Domestic animals | -76.2 | $6.41 \times 10^{6}$ | $-1.19 \times 10^{-5}$ | 1 |
| Farms | 33.5 | $2.78 \times 10^{6}$ | $1.2 \times 10^{-5}$ | 1 |
| Flora, fauna | -78.9 | $8.23 \times 10^{6}$ | $-9.59 \times 10^{-6}$ | 1 |
| Human samples | 34.6 | $2.78 \times 10^{6}$ | $1.24 \times 10^{-5}$ | 1 |
| Human habitat | -76.3 | $1.07 \times 10^{7}$ | $-7.11 \times 10^{-6}$ | 1 |
| Freshwater | -91.7 | $8.84 \times 10^{6}$ | $-1.04 \times 10^{-5}$ | 1 |
| Clinical samples | 34.9 | $2.78 \times 10^{6}$ | $1.25 \times 10^{-5}$ | 1 |
| Sludge, waste | -75.8 | $5.17 \times 10^{6}$ | $-1.46 \times 10^{-5}$ | 1 |
| Soil | -82.4 | $6.25 \times 10^{6}$ | $-1.32 \times 10^{-5}$ | 1 |

**Table S13: Summary of the selected model for CHG 14.** φ correspond to the estimated time autoregression coefficient. ν and ρ are respectively the estimated smoothness and scaling parameters of the Matérn correlation function. *: $p<0.05$, **: $p<0.01$, ***: $p<0.001$.

$\varphi = 5.13 \times 10^{-4}$, $\nu = 5.24 \times 10^{-3}$, $\rho = 0.724$

| **Variable** | **Estimate** | **Conditional standard error** | **t** | **p** |
|---|---|---|---|---|
| Intercept | -39 | $3.05 \times 10^{6}$ | $-1.28 \times 10^{-5}$ | 1 |
| Domestic animals | -16.3 | $6.53 \times 10^{6}$ | $-2.49 \times 10^{-6}$ | 1 |
| Farms | 33.8 | $3.05 \times 10^{6}$ | $1.11 \times 10^{-5}$ | 1 |
| Flora, fauna | -17 | $8.33 \times 10^{6}$ | $-2.04 \times 10^{-6}$ | 1 |
| Human samples | -16.1 | $3.6 \times 10^{6}$ | $-4.46 \times 10^{-6}$ | 1 |
| Human habitat | -17.9 | $1.08 \times 10^{7}$ | $-1.66 \times 10^{-6}$ | 1 |
| Freshwater | -15.8 | $8.93 \times 10^{6}$ | $-1.77 \times 10^{-6}$ | 1 |
| Clinical samples | 30.8 | $3.05 \times 10^{6}$ | $1.01 \times 10^{-5}$ | 1 |
| Sludge, waste | -18.1 | $5.32 \times 10^{6}$ | $-3.4 \times 10^{-6}$ | 1 |
| Soil | -16.8 | $6.37 \times 10^{6}$ | $-2.63 \times 10^{-6}$ | 1 |
| Trade | 0.893 | 0.135 | 6.6 | $4.19 \times 10^{-11}$ *** |
| Migration | 0.173 | 0.242 | 0.717 | 0.473 |

**Table S14: Summary of the selected model for CHG 17.1.** φ correspond to the estimated time autoregression coefficient. ν and ρ are respectively the estimated smoothness and scaling parameters of the Matérn correlation function. *: p<0.05, **: p<0.01, ***: p<0.001.

$\varphi = 4.14 \times 10^{-3}$, $\nu = 16.7$, $\rho = 0.379$

| **Variable** | **Estimate** | **Conditional standard error** | **t** | **p** |
|---|---|---|---|---|
| Intercept | -39.9 | $2.83 \times 10^{6}$ | $-1.41 \times 10^{-5}$ | 1 |
| Domestic animals | $-1.95 \times 10^{2}$ | $6.43 \times 10^{6}$ | $-3.03 \times 10^{-5}$ | 1 |
| Farms | $-1.97 \times 10^{2}$ | $3.2 \times 10^{6}$ | $-6.14 \times 10^{-5}$ | 1 |
| Flora, fauna | $-1.97 \times 10^{2}$ | $8.25 \times 10^{6}$ | $-2.39 \times 10^{-5}$ | 1 |
| Human samples | 29.2 | $2.83 \times 10^{6}$ | $1.03 \times 10^{-5}$ | 1 |
| Human habitat | $-1.97 \times 10^{2}$ | $1.07 \times 10^{7}$ | $-1.83 \times 10^{-5}$ | 1 |
| Freshwater | 31.7 | $2.83 \times 10^{6}$ | $1.12 \times 10^{-5}$ | 1 |
| Clinical samples | 31.5 | $2.83 \times 10^{6}$ | $1.11 \times 10^{-5}$ | 1 |
| Sludge, waste | 32 | $2.83 \times 10^{6}$ | $1.13 \times 10^{-5}$ | 1 |
| Soil | $-1.98 \times 10^{2}$ | $6.27 \times 10^{6}$ | $-3.15 \times 10^{-5}$ | 1 |
| Trade | 0.805 | 0.29 | 2.77 | $5.52 \times 10^{-3}$ ** |
| Migration | $8.64 \times 10^{-2}$ | 0.389 | 0.222 | 0.824 |
| Aminoglycosides | -3.95 | 0.782 | -5.05 | $4.43 \times 10^{-7}$ *** |

**Table S15: Summary of the selected model for CHG 19.** φ correspond to the estimated time autoregression coefficient. ν and ρ are respectively the estimated smoothness and scaling parameters of the Matérn correlation function. *: $p<0.05$, **: $p<0.01$, ***: $p<0.001$.

φ = 0.842, ν = 16.7, ρ = 3.23

| **Variable** | **Estimate** | **Conditional standard error** | **t** | **p** |
|---|---|---|---|---|
| Intercept | -6.95 | 1.38 | -5.03 | $4.87 \times 10^{-7}$ *** |
| Domestic animals | -15.6 | 11.9 | -1.31 | 0.19 |
| Farms | 1.32 | 1.27 | 1.04 | 0.297 |
| Flora, fauna | -2.06 | 5.33 | -0.386 | 0.699 |
| Human samples | 1.71 | 1.28 | 1.33 | 0.183 |
| Human habitat | $-1.33 \times 10^{2}$ | $1.98 \times 10^{6}$ | $-6.71 \times 10^{-5}$ | 1 |
| Freshwater | -30.1 | $8.69 \times 10^{6}$ | $-3.46 \times 10^{-6}$ | 1 |
| Clinical samples | 3.24 | 1.21 | 2.68 | $7.41 \times 10^{-3}$ ** |
| Sludge, waste | 2.1 | 1.49 | 1.41 | 0.159 |
| Soil | -29.8 | $7.58 \times 10^{6}$ | $-3.94 \times 10^{-6}$ | 1 |
| Trade | 0.169 | 0.697 | 0.243 | 0.808 |
| Migration | 0.997 | 1.27 | 0.785 | 0.433 |
| Aminoglycosides | -0.643 | 0.199 | -3.23 | $1.25 \times 10^{-3}$ ** |
| Domestic animals × Trade | -1.5 | 0.956 | -1.57 | 0.117 |
| Farms × Trade | 0.362 | 0.751 | 0.482 | 0.63 |
| Flora, fauna × Trade | -12.3 | 7.89 | -1.56 | 0.12 |
| Human samples × Trade | 0.329 | 0.721 | 0.456 | 0.648 |
| Human habitat × Trade | $1.17 \times 10^{2}$ | $1.39 \times 10^{6}$ | $8.39 \times 10^{-5}$ | 1 |
| Freshwater × Trade | -0.456 | $9.8 \times 10^{6}$ | $-4.65 \times 10^{-8}$ | 1 |
| Clinical samples × Trade | -0.497 | 0.697 | -0.713 | 0.476 |
| Sludge, waste × Trade | 0.779 | 0.885 | 0.881 | 0.378 |
| Soil × Trade | -0.613 | $7.35 \times 10^{6}$ | $-8.35 \times 10^{-8}$ | 1 |
| Domestic animals × Migration | 16.1 | 9.02 | 1.79 | $7.38 \times 10^{-2}$ * |
| Farms × Migration | -2.05 | 1.34 | -1.53 | 0.126 |
| Flora, fauna × Migration | -0.988 | 1.71 | -0.578 | 0.563 |
| Human samples × Migration | -0.633 | 1.32 | -0.48 | 0.631 |
| Human habitat × Migration | $-1.59 \times 10^{2}$ | $1.99 \times 10^{6}$ | $-7.98 \times 10^{-5}$ | 1 |
| Freshwater × Migration | -1.26 | $1.03 \times 10^{7}$ | $-1.22 \times 10^{-7}$ | 1 |
| Clinical samples × Migration | -1.02 | 1.26 | -0.809 | 0.419 |
| Sludge, waste × Migration | -0.713 | 1.44 | -0.493 | 0.622 |
| Soil × Migration | -1.08 | $7.66 \times 10^{6}$ | $-1.41 \times 10^{-7}$ | 1 |

**Table S16: Summary of the selected model for CHG 20.** φ correspond to the estimated time autoregression coefficient. ν and ρ are respectively the estimated smoothness and scaling parameters of the Matérn correlation function. *: p<0.05, **: p<0.01, ***: p<0.001.

φ = -0.922, ν = 0.65, ρ = 10.3

| **Variable** | **Estimate** | **Conditional standard error** | **t** | **p** |
|---|---|---|---|---|
| Intercept | -39.1 | $3.01 \times 10^{6}$ | $-1.3 \times 10^{-5}$ | 1 |
| Domestic animals | 1.73 | $6.89 \times 10^{6}$ | $2.51 \times 10^{-7}$ | 1 |
| Farms | 33.7 | $3.01 \times 10^{6}$ | $1.12 \times 10^{-5}$ | 1 |
| Flora, fauna | 1.58 | $9.19 \times 10^{6}$ | $1.71 \times 10^{-7}$ | 1 |
| Human samples | 2.48 | $3.64 \times 10^{6}$ | $6.81 \times 10^{-7}$ | 1 |
| Human habitat | 0.782 | $1.23 \times 10^{7}$ | $6.35 \times 10^{-8}$ | 1 |
| Freshwater | 1.97 | $9.04 \times 10^{6}$ | $2.18 \times 10^{-7}$ | 1 |
| Clinical samples | 30.9 | $3.01 \times 10^{6}$ | $1.03 \times 10^{-5}$ | 1 |
| Sludge, waste | -0.653 | $5.3 \times 10^{6}$ | $-1.23 \times 10^{-7}$ | 1 |
| Soil | 1.69 | $7.3 \times 10^{6}$ | $2.31 \times 10^{-7}$ | 1 |
| Trade | 1.04 | 0.198 | 5.26 | $1.44 \times 10^{-7}$ *** |
| Migration | 0.117 | 0.343 | 0.342 | 0.732 |

**Table S17: Summary of the selected model for CHG 23.** φ correspond to the estimated time autoregression coefficient. ν and ρ are respectively the estimated smoothness and scaling parameters of the Matérn correlation function. *: $p<0.05$, **: $p<0.01$, ***: $p<0.001$.

$\varphi = -0.738$, $\nu = 4.83 \times 10^{-2}$, $\rho = 6.65$

| Variable | Estimate | Conditional standard error | t | p |
|---|---|---|---|---|
| Intercept | -41.4 | $3.52 \times 10^{6}$ | $-1.17 \times 10^{-5}$ | 1 |
| Domestic animals | -8.59 | $6.76 \times 10^{6}$ | $-1.27 \times 10^{-6}$ | 1 |
| Farms | -13.3 | $3.82 \times 10^{6}$ | $-3.48 \times 10^{-6}$ | 1 |
| Flora, fauna | -16.6 | $8.51 \times 10^{6}$ | $-1.95 \times 10^{-6}$ | 1 |
| Human samples | 29.3 | $3.52 \times 10^{6}$ | $8.33 \times 10^{-6}$ | 1 |
| Human habitat | -16.2 | $1.09 \times 10^{7}$ | $-1.48 \times 10^{-6}$ | 1 |
| Freshwater | -37.4 | $9.1 \times 10^{6}$ | $-4.11 \times 10^{-6}$ | 1 |
| Clinical samples | 32.1 | $3.52 \times 10^{6}$ | $9.12 \times 10^{-6}$ | 1 |
| Sludge, waste | -11.9 | $5.6 \times 10^{6}$ | $-2.12 \times 10^{-6}$ | 1 |
| Soil | -21.5 | $6.61 \times 10^{6}$ | $-3.26 \times 10^{-6}$ | 1 |
| Trade | 1.12 | 0.441 | 2.54 | $1.12 \times 10^{-2}$ * |
| Migration | -0.395 | 0.283 | -1.4 | 0.163 |

**Table S18: Summary of the selected model for CHG 27.** φ correspond to the estimated time autoregression coefficient. ν and ρ are respectively the estimated smoothness and scaling parameters of the Matérn correlation function. *: $p<0.05$, **: $p<0.01$, ***: $p<0.001$.

$\varphi = 1.31 \times 10^{-7}$, $\nu = 0.33$, $\rho = 7.09 \times 10^{-2}$

| **Variable** | **Estimate** | **Conditional standard error** | **t** | **p** |
|---|---|---|---|---|
| Intercept | -7.25 | 2.95 | -2.46 | $1.39 \times 10^{-2}$ * |
| Domestic animals | -30.7 | $8.31 \times 10^{6}$ | $-3.69 \times 10^{-6}$ | 1 |
| Farms | -6.4 | 1.97 | -3.26 | $1.13 \times 10^{-3}$ ** |
| Flora, fauna | -31.2 | $1.54 \times 10^{7}$ | $-2.02 \times 10^{-6}$ | 1 |
| Human samples | -0.577 | 0.691 | -0.836 | 0.403 |
| Human habitat | -22.9 | $5.58 \times 10^{7}$ | $-4.09 \times 10^{-7}$ | 1 |
| Freshwater | -2.88 | 5.5 | -0.524 | 0.6 |
| Clinical samples | -0.555 | 0.478 | -1.16 | 0.246 |
| Sludge, waste | -30 | $4.52 \times 10^{6}$ | $-6.64 \times 10^{-6}$ | 1 |
| Soil | -0.701 | 1.04 | -0.676 | 0.499 |
| Trade | -1.67 | 0.216 | -7.75 | $9.1 \times 10^{-15}$ *** |
| Migration | -2.77 | 0.74 | -3.74 | $1.85 \times 10^{-4}$ *** |
| Domestic animals × Trade | -1 | $9.69 \times 10^{6}$ | $-1.03 \times 10^{-7}$ | 1 |
| Farms × Trade | 3.06 | 0.886 | 3.46 | $5.44 \times 10^{-4}$ *** |
| Flora, fauna × Trade | -0.336 | $8.97 \times 10^{6}$ | $-3.74 \times 10^{-8}$ | 1 |
| Human samples × Trade | 0.183 | 0.914 | 0.2 | 0.841 |
| Human habitat × Trade | -5.51 | $5.58 \times 10^{7}$ | $-9.87 \times 10^{-8}$ | 1 |
| Freshwater × Trade | -4.57 | 11.2 | -0.408 | 0.683 |
| Clinical samples × Trade | -1.17 | 0.64 | -1.83 | $6.69 \times 10^{-2}$ * |
| Sludge, waste × Trade | -0.979 | $5.62 \times 10^{6}$ | $-1.74 \times 10^{-7}$ | 1 |
| Soil × Trade | 0.215 | 0.642 | 0.335 | 0.737 |
| Domestic animals × Migration | -1.04 | $1.34 \times 10^{7}$ | $-7.8 \times 10^{-8}$ | 1 |
| Farms × Migration | -10.1 | 2.73 | -3.68 | $2.31 \times 10^{-4}$ *** |
| Flora, fauna × Migration | -1.17 | $2.83 \times 10^{7}$ | $-4.13 \times 10^{-8}$ | 1 |
| Human samples × Migration | -2.9 | 0.894 | -3.25 | $1.16 \times 10^{-3}$ ** |
| Human habitat × Migration | 13 | $1.06 \times 10^{8}$ | $1.22 \times 10^{-7}$ | 1 |
| Freshwater × Migration | -4.33 | 7.94 | -0.546 | 0.585 |
| Clinical samples × Migration | -2.16 | 0.334 | -6.48 | $9.4 \times 10^{-11}$ *** |
| Sludge, waste × Migration | -2.77 | $5.52 \times 10^{6}$ | $-5.01 \times 10^{-7}$ | 1 |
| Soil × Migration | -1.46 | 0.87 | -1.68 | $9.39 \times 10^{-2}$ * |

**Table S19: Human exchanges variables influencing the prevalence of ARBs**

| | CHG 2 | CHG 3 | CHG 4 | CHG 5.1 | CHG 6.1 | CHG 7 | CHG 8 | CHG 11 | CHG 14 | CHG 17.1 | CHG 19 | CHG 20 | CHG 23 | CHG 27 |
|---|---|---|---|---|---|---|---|---|---|---|---|---|---|---|
| Animal and vegetable oils, fats and waxes | | | | | | | | | × | × | | | × | |
| Beverages | × | | | | | | | | × | × | | × | × | |
| Cereals and cereal preparations | | × | | | | | | | × | × | | × | | |
| Coal, coke and briquettes | | | | | | | | | × | × | | × | | |
| Coffee, tea, cocoa, spices, and manufactures thereof | | | | | | | | | × | × | | | | |
| Cork and wood | | | × | | | | | | × | × | | × | | |
| Crude materials, inedible, except fuels | | | | | | | | | | × | | | | |
| Crude rubber (including synthetic and reclaimed) | | | | × | | | | | | × | | | × | |
| Dairy products and birds' eggs | | | | | | | | | × | × | | × | × | |
| Dyeing, tanning and colouring materials | | × | | | | | | | × | × | | | × | |
| Feedstuff for animals (excluding unmilled cereals) | × | | | | | | | | × | × | | × | | |
| Fish, crustaceans, molluscs and preparations thereof | | | | | | | | | × | × | | × | | |
| Food and live animals | | | | | | | | | × | × | | × | | |
| Hides, skins and furskins, raw | | | | | | | | | | × | | | × | |
| Live animals other than animals of division 03 | | × | | | | | | | | × | | | | |
| Manufactured goods | | | | | | | | | × | × | | × | | |
| Meat and meat preparations | × | | | | | | | | × | × | × | × | | |
| Medicinal and pharmaceutical products | × | | | | | | | | × | × | | | × | |
| Mineral fuels, lubricants and related materials | × | | | | | | | × | × | × | | | | |
| Miscellaneous edible products and preparations | × | | | | | | | | × | × | | | × | |
| Oil seeds and oleaginous fruits | | × | | | × | × | | | | × | | | × | |
| Petroleum, petroleum products and related materials | × | | | | | | | | × | × | | | | |
| Plastics | | | | | | | | | × | × | | | × | |
| Sugar, sugar preparations and honey | × | | | | | | × | | × | × | × | | | |
| Textiles fibres and their wastes | | | × | × | | | | | | × | | | | |
| Vegetables and fruits | | | | | | | | | × | × | | × | | |
| Human migration | × | | | | | | × | | | × | | | | × |

**Table S20: Summary of the selected model for CHG 1 for worldwide data.** φ correspond to the estimated time autoregression coefficient. ν and ρ are respectively the estimated smoothness and scaling parameters of the Matérn correlation function. *: $p<0.05$, **: $p<0.01$, ***: $p<0.001$.

$\varphi = 0.264$, $\nu = 16.7$, $\rho = 0.729$

| **Variable** | **Estimate** | **Conditional standard error** | **t** | **p** |
|---|---|---|---|---|
| Intercept | -7.91 | 0.833 | -9.51 | 0 *** |
| Domestic animals | -29.8 | $4.08 \times 10^{6}$ | $-7.32 \times 10^{-6}$ | 1 |
| Farms | 0.411 | 0.76 | 0.541 | 0.589 |
| Flora, fauna | -29.9 | $3.97 \times 10^{6}$ | $-7.54 \times 10^{-6}$ | 1 |
| Human samples | -0.307 | 0.815 | -0.376 | 0.707 |
| Human habitat | 0.279 | 1.24 | 0.225 | 0.822 |
| Freshwater | -0.622 | 1.23 | -0.504 | 0.614 |
| Clinical samples | 1.65 | 0.721 | 2.28 | $2.23 \times 10^{-2}$ * |
| Sludge, waste | 2.2 | 0.933 | 2.35 | $1.86 \times 10^{-2}$ * |
| Soil | -30.1 | $2.02 \times 10^{6}$ | $-1.49 \times 10^{-5}$ | 1 |
| Trade | -0.136 | $7.04 \times 10^{-2}$ | -1.93 | $5.36 \times 10^{-2}$ * |
| Migration | -0.162 | $7.78 \times 10^{-2}$ | -2.09 | $3.68 \times 10^{-2}$ * |

**Table S21: Summary of the selected model for CHG 2 for worldwide data.** φ correspond to the estimated time autoregression coefficient. ν and ρ are respectively the estimated smoothness and scaling parameters of the Matérn correlation function. *: $p<0.05$, **: $p<0.01$, ***: $p<0.001$.

$\varphi = -0.167$, $\nu = 9.1 \times 10^{-3}$, $\rho = 1.48$

| **Variable** | **Estimate** | **Conditional standard error** | **t** | **p** |
|---|---|---|---|---|
| Intercept | -5.81 | 0.556 | -10.4 | 0 *** |
| Domestic animals | -32 | $4.43 \times 10^{6}$ | $-7.22 \times 10^{-6}$ | 1 |
| Farms | -0.117 | 0.277 | -0.423 | 0.673 |
| Flora, fauna | -20.2 | 23.9 | -0.846 | 0.397 |
| Human samples | -2.84 | 0.657 | -4.32 | $1.57 \times 10^{-5}$ *** |
| Human habitat | -31.6 | $4.08 \times 10^{6}$ | $-7.74 \times 10^{-6}$ | 1 |
| Freshwater | -3.6 | 1.68 | -2.15 | $3.17 \times 10^{-2}$ * |
| Clinical samples | -2.04 | 0.282 | -7.26 | $3.98 \times 10^{-13}$ *** |
| Sludge, waste | -31.1 | $3.88 \times 10^{6}$ | $-8 \times 10^{-6}$ | 1 |
| Soil | -1.4 | 0.588 | -2.38 | $1.73 \times 10^{-2}$ * |
| Trade | -1.03 | 0.317 | -3.26 | $1.11 \times 10^{-3}$ ** |
| Migration | 1.09 | 0.168 | 6.5 | $8.08 \times 10^{-11}$ *** |
| Domestic animals × Trade | 0.743 | $2.22 \times 10^{6}$ | $3.35 \times 10^{-7}$ | 1 |
| Farms × Trade | 0.95 | 0.307 | 3.1 | $1.97 \times 10^{-3}$ ** |
| Flora, fauna × Trade | -83.2 | 87.9 | -0.947 | 0.343 |
| Human samples × Trade | -0.455 | 1.78 | -0.256 | 0.798 |
| Human habitat × Trade | 0.741 | $1.17 \times 10^{6}$ | $6.33 \times 10^{-7}$ | 1 |
| Freshwater × Trade | -7.78 | 5.67 | -1.37 | 0.17 |
| Clinical samples × Trade | 0.958 | 0.31 | 3.09 | $2.03 \times 10^{-3}$ ** |
| Sludge, waste × Trade | 0.618 | $3.11 \times 10^{6}$ | $1.99 \times 10^{-7}$ | 1 |
| Soil × Trade | 0.781 | 0.566 | 1.38 | 0.167 |
| Domestic animals × Migration | -1.01 | $2.19 \times 10^{6}$ | $-4.62 \times 10^{-7}$ | 1 |
| Farms × Migration | -1.33 | 0.216 | -6.15 | $7.85 \times 10^{-10}$ *** |
| Flora, fauna × Migration | -0.18 | 2.61 | $-6.88 \times 10^{-2}$ | 0.945 |
| Human samples × Migration | -2.09 | 1.46 | -1.43 | 0.154 |
| Human habitat × Migration | -1.46 | $4.82 \times 10^{6}$ | $-3.03 \times 10^{-7}$ | 1 |
| Freshwater × Migration | -3.82 | 4.09 | -0.932 | 0.351 |
| Clinical samples × Migration | -1.61 | 0.313 | -5.13 | $2.84 \times 10^{-7}$ *** |
| Sludge, waste × Migration | -1.28 | $3.76 \times 10^{6}$ | $-3.41 \times 10^{-7}$ | 1 |
| Soil × Migration | -1.21 | 0.856 | -1.41 | 0.158 |

**Table S22: Summary of the selected model for CHG 3 for worldwide data.** φ correspond to the estimated time autoregression coefficient. ν and ρ are respectively the estimated smoothness and scaling parameters of the Matérn correlation function. *: $p<0.05$, **: $p<0.01$, ***: $p<0.001$.

$\varphi = -0.208$, $\nu = 1.53$, $\rho = 2.18$

| **Variable** | **Estimate** | **Conditional standard error** | **t** | **p** |
|---|---|---|---|---|
| Intercept | -8.37 | 0.984 | -8.51 | 0 *** |
| Domestic animals | $-1.94 \times 10^{3}$ | $2.9 \times 10^{5}$ | $-6.71 \times 10^{-3}$ | 0.995 |
| Farms | -1.59 | 1.44 | -1.1 | 0.27 |
| Flora, fauna | 2.83 | 1.12 | 2.54 | $1.12 \times 10^{-2}$ * |
| Human samples | 2.92 | 0.857 | 3.4 | $6.64 \times 10^{-4}$ *** |
| Human habitat | 0.778 | 1.57 | 0.495 | 0.621 |
| Freshwater | $3.3 \times 10^{-2}$ | 1.15 | $2.85 \times 10^{-2}$ | 0.977 |
| Clinical samples | 3.48 | 0.853 | 4.08 | $4.59 \times 10^{-5}$ *** |
| Sludge, waste | -1.08 | 2.33 | -0.464 | 0.643 |
| Soil | 2.36 | 0.888 | 2.66 | $7.88 \times 10^{-3}$ ** |
| Trade | -0.7 | 0.429 | -1.63 | 0.103 |
| Migration | -1.41 | 1.63 | -0.864 | 0.387 |
| Domestic animals × Trade | $2.14 \times 10^{2}$ | $3.2 \times 10^{4}$ | $6.67 \times 10^{-3}$ | 0.995 |
| Farms × Trade | -3.17 | 2.17 | -1.46 | 0.143 |
| Flora, fauna × Trade | -1.18 | 1.49 | -0.793 | 0.428 |
| Human samples × Trade | -0.76 | 0.449 | -1.69 | $9.08 \times 10^{-2}$ * |
| Human habitat × Trade | $9.91 \times 10^{-2}$ | 0.992 | 0.1 | 0.92 |
| Freshwater × Trade | -0.7 | 0.512 | -1.37 | 0.172 |
| Clinical samples × Trade | 0.495 | 0.429 | 1.15 | 0.249 |
| Sludge, waste × Trade | -4.15 | 3.48 | -1.19 | 0.233 |
| Soil × Trade | 0.743 | 0.453 | 1.64 | 0.1 |
| Domestic animals × Migration | $-2.94 \times 10^{3}$ | $4.37 \times 10^{5}$ | $-6.72 \times 10^{-3}$ | 0.995 |
| Farms × Migration | -0.703 | 2.21 | -0.318 | 0.75 |
| Flora, fauna × Migration | 0.675 | 1.72 | 0.392 | 0.695 |
| Human samples × Migration | 1.55 | 1.63 | 0.95 | 0.342 |
| Human habitat × Migration | $-5.3 \times 10^{-2}$ | 2.47 | $-2.14 \times 10^{-2}$ | 0.983 |
| Freshwater × Migration | -6.51 | 2.14 | -3.04 | $2.4 \times 10^{-3}$ ** |
| Clinical samples × Migration | 1.44 | 1.63 | 0.882 | 0.378 |
| Sludge, waste × Migration | -4.59 | 5.36 | -0.857 | 0.391 |
| Soil × Migration | $7.08 \times 10^{-2}$ | 1.67 | $4.25 \times 10^{-2}$ | 0.966 |

**Table S23: Summary of the selected model for CHG 4 for worldwide data.** φ correspond to the estimated time autoregression coefficient. ν and ρ are respectively the estimated smoothness and scaling parameters of the Matérn correlation function. *: $p<0.05$, **: $p<0.01$, ***: $p<0.001$.

$\varphi = 0.216$, $\nu = 5.49 \times 10^{-3}$, $\rho = 2.12$

| **Variable** | **Estimate** | **Conditional standard error** | **t** | **p** |
|---|---|---|---|---|
| Intercept | -6.84 | 0.698 | -9.81 | 0 *** |
| Domestic animals | -30.7 | $4.59 \times 10^{6}$ | $-6.69 \times 10^{-6}$ | 1 |
| Farms | 0.109 | 0.49 | 0.222 | 0.824 |
| Flora, fauna | 2.27 | 0.598 | 3.79 | $1.48 \times 10^{-4}$ *** |
| Human samples | -1.99 | 0.956 | -2.08 | $3.71 \times 10^{-2}$ * |
| Human habitat | -3.12 | 1.62 | -1.93 | $5.37 \times 10^{-2}$ * |
| Freshwater | -3.28 | 3.4 | -0.965 | 0.335 |
| Clinical samples | -0.106 | 0.459 | -0.231 | 0.817 |
| Sludge, waste | -31.2 | $5.99 \times 10^{6}$ | $-5.21 \times 10^{-6}$ | 1 |
| Soil | -1.45 | 1.24 | -1.17 | 0.243 |
| Trade | 0.328 | 0.437 | 0.752 | 0.452 |
| Migration | -1.8 | 0.983 | -1.83 | $6.7 \times 10^{-2}$ * |
| Domestic animals × Trade | -1.12 | $6.93 \times 10^{6}$ | $-1.62 \times 10^{-7}$ | 1 |
| Farms × Trade | -1.59 | 0.527 | -3.01 | $2.6 \times 10^{-3}$ ** |
| Flora, fauna × Trade | -2.47 | 1.16 | -2.13 | $3.32 \times 10^{-2}$ * |
| Human samples × Trade | -1.03 | 0.741 | -1.39 | 0.163 |
| Human habitat × Trade | 0.215 | 0.973 | 0.221 | 0.825 |
| Freshwater × Trade | -16.4 | 14.4 | -1.14 | 0.254 |
| Clinical samples × Trade | -0.478 | 0.44 | -1.09 | 0.277 |
| Sludge, waste × Trade | -0.682 | $5.94 \times 10^{6}$ | $-1.15 \times 10^{-7}$ | 1 |
| Soil × Trade | -0.349 | 1.21 | -0.288 | 0.774 |
| Domestic animals × Migration | 2.23 | $7.94 \times 10^{6}$ | $2.81 \times 10^{-7}$ | 1 |
| Farms × Migration | 0.544 | 1.11 | 0.488 | 0.625 |
| Flora, fauna × Migration | 1.62 | 1.24 | 1.31 | 0.191 |
| Human samples × Migration | -3.29 | 3.33 | -0.986 | 0.324 |
| Human habitat × Migration | -9.21 | 6.46 | -1.43 | 0.154 |
| Freshwater × Migration | -1.16 | 6.12 | -0.189 | 0.85 |
| Clinical samples × Migration | 1.42 | 0.985 | 1.44 | 0.15 |
| Sludge, waste × Migration | 1.82 | $6.44 \times 10^{6}$ | $2.83 \times 10^{-7}$ | 1 |
| Soil × Migration | 1.22 | 1.8 | 0.68 | 0.496 |

**Table S24: Summary of the selected model for CHG 5.1 for worldwide data.** φ correspond to the estimated time autoregression coefficient. ν and ρ are respectively the estimated smoothness and scaling parameters of the Matérn correlation function. *: p<0.05, **: p<0.01, ***: p<0.001.

$\varphi = 0.755$, $\nu = 1.27 \times 10^{-2}$, $\rho = 1.17$

| **Variable** | **Estimate** | **Conditional standard error** | **t** | **p** |
|---|---|---|---|---|
| Intercept | $-1.43 \times 10^{2}$ | $6.01 \times 10^{3}$ | $-2.38 \times 10^{-2}$ | 0.981 |
| Domestic animals | $1.33 \times 10^{2}$ | $6.01 \times 10^{3}$ | $2.22 \times 10^{-2}$ | 0.982 |
| Farms | $1.35 \times 10^{2}$ | $6.01 \times 10^{3}$ | $2.25 \times 10^{-2}$ | 0.982 |
| Flora, fauna | $1.38 \times 10^{2}$ | $6.01 \times 10^{3}$ | $2.3 \times 10^{-2}$ | 0.982 |
| Human samples | $1.37 \times 10^{2}$ | $6.01 \times 10^{3}$ | $2.28 \times 10^{-2}$ | 0.982 |
| Human habitat | $1.37 \times 10^{2}$ | $6.01 \times 10^{3}$ | $2.29 \times 10^{-2}$ | 0.982 |
| Freshwater | $-1.55 \times 10^{2}$ | $6.83 \times 10^{4}$ | $-2.28 \times 10^{-3}$ | 0.998 |
| Clinical samples | $1.39 \times 10^{2}$ | $6.01 \times 10^{3}$ | $2.31 \times 10^{-2}$ | 0.982 |
| Sludge, waste | $1.39 \times 10^{2}$ | $6.01 \times 10^{3}$ | $2.31 \times 10^{-2}$ | 0.982 |
| Soil | $1.32 \times 10^{2}$ | $6.01 \times 10^{3}$ | $2.2 \times 10^{-2}$ | 0.982 |
| Trade | 32.5 | $1.43 \times 10^{3}$ | $2.27 \times 10^{-2}$ | 0.982 |
| Migration | -7.17 | $3.77 \times 10^{2}$ | $-1.9 \times 10^{-2}$ | 0.985 |
| Domestic animals × Trade | -31.3 | $1.43 \times 10^{3}$ | $-2.18 \times 10^{-2}$ | 0.983 |
| Farms × Trade | -31.8 | $1.43 \times 10^{3}$ | $-2.22 \times 10^{-2}$ | 0.982 |
| Flora, fauna × Trade | -33.2 | $1.43 \times 10^{3}$ | $-2.32 \times 10^{-2}$ | 0.982 |
| Human samples × Trade | -31.7 | $1.43 \times 10^{3}$ | $-2.21 \times 10^{-2}$ | 0.982 |
| Human habitat × Trade | -32.6 | $1.43 \times 10^{3}$ | $-2.28 \times 10^{-2}$ | 0.982 |
| Freshwater × Trade | 33.7 | $1.54 \times 10^{4}$ | $2.19 \times 10^{-3}$ | 0.998 |
| Clinical samples × Trade | -32.9 | $1.43 \times 10^{3}$ | $-2.29 \times 10^{-2}$ | 0.982 |
| Sludge, waste × Trade | -32.6 | $1.43 \times 10^{3}$ | $-2.27 \times 10^{-2}$ | 0.982 |
| Soil × Trade | -32 | $1.43 \times 10^{3}$ | $-2.24 \times 10^{-2}$ | 0.982 |
| Domestic animals × Migration | 6.77 | $3.77 \times 10^{2}$ | $1.8 \times 10^{-2}$ | 0.986 |
| Farms × Migration | 6.55 | $3.77 \times 10^{2}$ | $1.74 \times 10^{-2}$ | 0.986 |
| Flora, fauna × Migration | 6.58 | $3.77 \times 10^{2}$ | $1.75 \times 10^{-2}$ | 0.986 |
| Human samples × Migration | 6.48 | $3.77 \times 10^{2}$ | $1.72 \times 10^{-2}$ | 0.986 |
| Human habitat × Migration | 6.99 | $3.77 \times 10^{2}$ | $1.85 \times 10^{-2}$ | 0.985 |
| Freshwater × Migration | $-4.24 \times 10^{2}$ | $1.01 \times 10^{5}$ | $-4.18 \times 10^{-3}$ | 0.997 |
| Clinical samples × Migration | 7.33 | $3.77 \times 10^{2}$ | $1.94 \times 10^{-2}$ | 0.984 |
| Sludge, waste × Migration | 6.91 | $3.77 \times 10^{2}$ | $1.83 \times 10^{-2}$ | 0.985 |
| Soil × Migration | 1.92 | $3.77 \times 10^{2}$ | $5.09 \times 10^{-3}$ | 0.996 |

**Table S25: Summary of the selected model for CHG 6.1 for worldwide data.** φ correspond to the estimated time autoregression coefficient. ν and ρ are respectively the estimated smoothness and scaling parameters of the Matérn correlation function. *: p<0.05, **: p<0.01, ***: p<0.001.

$\varphi = 0.747$, $\nu = 0.415$, $\rho = 4.37 \times 10^{-2}$

| **Variable** | **Estimate** | **Conditional standard error** | **t** | **p** |
|---|---|---|---|---|
| Intercept | -3.24 | 0.718 | -4.51 | $6.49 \times 10^{-6}$ *** |
| Domestic animals | -1.2 | 0.417 | -2.88 | $3.94 \times 10^{-3}$ ** |
| Farms | 1.03 | 0.134 | 7.71 | $1.28 \times 10^{-14}$ *** |
| Flora, fauna | 1.31 | 0.247 | 5.3 | $1.13 \times 10^{-7}$ *** |
| Human samples | 0.5 | 0.133 | 3.75 | $1.77 \times 10^{-4}$ *** |
| Human habitat | 0.329 | 0.269 | 1.22 | 0.221 |
| Freshwater | -0.541 | 0.287 | -1.88 | $5.96 \times 10^{-2}$ * |
| Clinical samples | -1.25 | 0.133 | -9.34 | 0 *** |
| Sludge, waste | -0.25 | 0.419 | -0.597 | 0.551 |
| Soil | -1.56 | 0.2 | -7.81 | $5.55 \times 10^{-15}$ *** |
| Trade | -0.131 | $8.09 \times 10^{-2}$ | -1.62 | 0.105 |
| Migration | -0.679 | 0.121 | -5.63 | $1.77 \times 10^{-8}$ *** |
| Domestic animals × Trade | -0.293 | 0.326 | -0.897 | 0.37 |
| Farms × Trade | -0.531 | $8.33 \times 10^{-2}$ | -6.38 | $1.73 \times 10^{-10}$ *** |
| Flora, fauna × Trade | -2.51 | 0.492 | -5.09 | $3.5 \times 10^{-7}$ *** |
| Human samples × Trade | -0.981 | $9.27 \times 10^{-2}$ | -10.6 | 0 *** |
| Human habitat × Trade | -0.654 | 0.227 | -2.87 | $4.04 \times 10^{-3}$ ** |
| Freshwater × Trade | -0.783 | 0.131 | -5.96 | $2.6 \times 10^{-9}$ *** |
| Clinical samples × Trade | -0.224 | $8.33 \times 10^{-2}$ | -2.68 | $7.3 \times 10^{-3}$ ** |
| Sludge, waste × Trade | -0.622 | 0.415 | -1.5 | 0.134 |
| Soil × Trade | 0.19 | 0.128 | 1.48 | 0.14 |
| Domestic animals × Migration | 0.102 | 0.342 | 0.3 | 0.764 |
| Farms × Migration | 0.179 | 0.116 | 1.55 | 0.121 |
| Flora, fauna × Migration | 0.26 | 0.25 | 1.04 | 0.299 |
| Human samples × Migration | 0.867 | 0.116 | 7.47 | $8.17 \times 10^{-14}$ *** |
| Human habitat × Migration | 0.833 | 0.222 | 3.76 | $1.72 \times 10^{-4}$ *** |
| Freshwater × Migration | -3.19 | 0.416 | -7.68 | $1.63 \times 10^{-14}$ *** |
| Clinical samples × Migration | 0.362 | 0.114 | 3.18 | $1.49 \times 10^{-3}$ ** |
| Sludge, waste × Migration | 0.692 | 0.251 | 2.76 | $5.72 \times 10^{-3}$ ** |
| Soil × Migration | -0.409 | 0.239 | -1.71 | $8.77 \times 10^{-2}$ * |

**Table S26: Summary of the selected model for CHG 7 for worldwide data.** φ correspond to the estimated time autoregression coefficient. ν and ρ are respectively the estimated smoothness and scaling parameters of the Matérn correlation function. *: $p<0.05$, **: $p<0.01$, ***: $p<0.001$.

$\varphi = -0.599$, $\nu = 16.7$, $\rho = 1.51$

| **Variable** | **Estimate** | **Conditional standard error** | **t** | **p** |
|---|---|---|---|---|
| Intercept | -4.9 | 0.437 | -11.2 | 0 *** |
| Domestic animals | 0.697 | 0.575 | 1.21 | 0.225 |
| Farms | 0.253 | 0.308 | 0.822 | 0.411 |
| Flora, fauna | 1.23 | 0.487 | 2.52 | $1.17 \times 10^{-2}$ * |
| Human samples | 0.697 | 0.292 | 2.38 | $1.71 \times 10^{-2}$ * |
| Human habitat | 0.398 | 0.501 | 0.796 | 0.426 |
| Freshwater | -0.776 | 0.592 | -1.31 | 0.19 |
| Clinical samples | 0.785 | 0.278 | 2.83 | $4.71 \times 10^{-3}$ ** |
| Sludge, waste | -1.03 | 1.6 | -0.642 | 0.521 |
| Soil | 0.321 | 0.451 | 0.711 | 0.477 |
| Trade | -0.111 | 0.257 | -0.434 | 0.664 |
| Migration | 0.182 | 0.145 | 1.25 | 0.21 |
| Domestic animals × Trade | $6.37 \times 10^{-2}$ | 0.43 | 0.148 | 0.882 |
| Farms × Trade | 0.229 | 0.267 | 0.857 | 0.391 |
| Flora, fauna × Trade | -1.88 | 1.16 | -1.62 | 0.105 |
| Human samples × Trade | 0.554 | 0.258 | 2.15 | $3.17 \times 10^{-2}$ * |
| Human habitat × Trade | -0.434 | 0.756 | -0.575 | 0.565 |
| Freshwater × Trade | -0.488 | 0.884 | -0.552 | 0.581 |
| Clinical samples × Trade | $-2.93 \times 10^{-3}$ | 0.259 | $-1.13 \times 10^{-2}$ | 0.991 |
| Sludge, waste × Trade | -3.69 | 4.33 | -0.852 | 0.394 |
| Soil × Trade | 0.593 | 0.287 | 2.06 | $3.9 \times 10^{-2}$ * |
| Domestic animals × Migration | -0.343 | 0.268 | -1.28 | 0.2 |
| Farms × Migration | -0.714 | 0.188 | -3.79 | $1.5 \times 10^{-4}$ *** |
| Flora, fauna × Migration | -0.519 | 0.578 | -0.897 | 0.37 |
| Human samples × Migration | -0.383 | 0.154 | -2.49 | $1.28 \times 10^{-2}$ * |
| Human habitat × Migration | 0.946 | 0.479 | 1.98 | $4.83 \times 10^{-2}$ * |
| Freshwater × Migration | -0.174 | 0.657 | -0.265 | 0.791 |
| Clinical samples × Migration | -0.376 | 0.147 | -2.55 | $1.09 \times 10^{-2}$ * |
| Sludge, waste × Migration | $1.43 \times 10^{-2}$ | 1.77 | $8.05 \times 10^{-3}$ | 0.994 |
| Soil × Migration | -1.93 | 0.747 | -2.58 | $9.84 \times 10^{-3}$ ** |

**Table S27: Summary of the selected model for CHG 8 for worldwide data.** φ correspond to the estimated time autoregression coefficient. ν and ρ are respectively the estimated smoothness and scaling parameters of the Matérn correlation function. *: p<0.05, **: p<0.01, ***: p<0.001.

φ = 0.28, ν = 16.7, ρ = 0.456

| **Variable** | **Estimate** | **Conditional standard error** | **t** | **p** |
|---|---|---|---|---|
| Intercept | -5.16 | 0.519 | -9.94 | 0 *** |
| Domestic animals | 0.615 | 0.519 | 1.18 | 0.236 |
| Farms | 0.941 | 0.274 | 3.43 | $5.98 \times 10^{-4}$ *** |
| Flora, fauna | 0.265 | 0.538 | 0.493 | 0.622 |
| Human samples | $-2.76 \times 10^{-2}$ | 0.28 | $-9.88 \times 10^{-2}$ | 0.921 |
| Human habitat | 0.471 | 0.396 | 1.19 | 0.234 |
| Freshwater | 0.299 | 0.569 | 0.526 | 0.599 |
| Clinical samples | 0.303 | 0.264 | 1.15 | 0.251 |
| Sludge, waste | -3.42 | 2.59 | -1.32 | 0.186 |
| Soil | 0.759 | 0.332 | 2.29 | $2.22 \times 10^{-2}$ * |
| Trade | $-4.04 \times 10^{-3}$ | 0.223 | $-1.81 \times 10^{-2}$ | 0.986 |
| Migration | -0.106 | 0.221 | -0.48 | 0.631 |
| Domestic animals × Trade | -0.799 | 0.616 | -1.3 | 0.195 |
| Farms × Trade | -0.482 | 0.24 | -2.01 | $4.45 \times 10^{-2}$ * |
| Flora, fauna × Trade | $3.12 \times 10^{-2}$ | 0.503 | $6.21 \times 10^{-2}$ | 0.95 |
| Human samples × Trade | 0.299 | 0.227 | 1.32 | 0.188 |
| Human habitat × Trade | 0.634 | 0.275 | 2.31 | $2.12 \times 10^{-2}$ * |
| Freshwater × Trade | -0.781 | 0.401 | -1.95 | $5.14 \times 10^{-2}$ * |
| Clinical samples × Trade | -0.201 | 0.225 | -0.895 | 0.371 |
| Sludge, waste × Trade | 0.237 | 1.39 | 0.171 | 0.864 |
| Soil × Trade | $2.5 \times 10^{-2}$ | 0.277 | $9.03 \times 10^{-2}$ | 0.928 |
| Domestic animals × Migration | -0.199 | 0.417 | -0.478 | 0.633 |
| Farms × Migration | -0.929 | 0.251 | -3.7 | $2.17 \times 10^{-4}$ *** |
| Flora, fauna × Migration | $-6.91 \times 10^{-2}$ | 0.428 | -0.161 | 0.872 |
| Human samples × Migration | $-5.02 \times 10^{-2}$ | 0.223 | -0.225 | 0.822 |
| Human habitat × Migration | -0.292 | 0.278 | -1.05 | 0.293 |
| Freshwater × Migration | -2.76 | 1.03 | -2.68 | $7.42 \times 10^{-3}$ ** |
| Clinical samples × Migration | -0.371 | 0.22 | -1.69 | $9.12 \times 10^{-2}$ * |
| Sludge, waste × Migration | 0.144 | 0.992 | 0.145 | 0.885 |
| Soil × Migration | -0.265 | 0.285 | -0.928 | 0.353 |

**Table S28: Summary of the selected model for CHG 11 for worldwide data.** φ correspond to the estimated time autoregression coefficient. ν and ρ are respectively the estimated smoothness and scaling parameters of the Matérn correlation function. *: p<0.05, **: p<0.01, ***: p<0.001.

φ = 0.671, ν = 16.7, ρ = 0.345

| Variable | Estimate | Conditional standard error | t | p |
|---|---|---|---|---|
| Intercept | -10.1 | 2.03 | -4.96 | $7.17 \times 10^{-7}$ *** |
| Domestic animals | -2.74 | 4.7 | -0.581 | 0.561 |
| Farms | 2.9 | 1.84 | 1.57 | 0.116 |
| Flora, fauna | 1.18 | 2.48 | 0.477 | 0.633 |
| Human samples | 3.56 | 1.84 | 1.94 | $5.27 \times 10^{-2}$ * |
| Human habitat | -16.9 | 17.1 | -0.99 | 0.322 |
| Freshwater | 1.77 | 1.95 | 0.908 | 0.364 |
| Clinical samples | 3.51 | 1.84 | 1.91 | $5.62 \times 10^{-2}$ * |
| Sludge, waste | -28 | $5.67 \times 10^{6}$ | $-4.93 \times 10^{-6}$ | 1 |
| Soil | 2.39 | 1.9 | 1.26 | 0.208 |
| Trade | -0.168 | 0.4 | -0.42 | 0.675 |
| Migration | -4.12 | 4.49 | -0.918 | 0.359 |
| Domestic animals × Trade | -7.07 | 5.47 | -1.29 | 0.196 |
| Farms × Trade | -0.644 | 0.429 | -1.5 | 0.133 |
| Flora, fauna × Trade | 0.665 | 0.764 | 0.871 | 0.384 |
| Human samples × Trade | 0.281 | 0.4 | 0.702 | 0.482 |
| Human habitat × Trade | -2.31 | 5.16 | -0.448 | 0.654 |
| Freshwater × Trade | -0.69 | 0.736 | -0.937 | 0.349 |
| Clinical samples × Trade | $-8.78 \times 10^{-2}$ | 0.399 | -0.22 | 0.826 |
| Sludge, waste × Trade | -0.4 | $5.77 \times 10^{6}$ | $-6.94 \times 10^{-8}$ | 1 |
| Soil × Trade | $7.43 \times 10^{-2}$ | 0.453 | 0.164 | 0.87 |
| Domestic animals × Migration | -5.16 | 12.2 | -0.422 | 0.673 |
| Farms × Migration | 4.43 | 4.49 | 0.986 | 0.324 |
| Flora, fauna × Migration | 2.4 | 5.45 | 0.441 | 0.659 |
| Human samples × Migration | 4.18 | 4.49 | 0.93 | 0.352 |
| Human habitat × Migration | -35.7 | 35.5 | -1.01 | 0.315 |
| Freshwater × Migration | 4.95 | 4.52 | 1.09 | 0.274 |
| Clinical samples × Migration | 3.65 | 4.49 | 0.814 | 0.416 |
| Sludge, waste × Migration | 4.59 | $7.67 \times 10^{6}$ | $5.98 \times 10^{-7}$ | 1 |
| Soil × Migration | 3.66 | 4.58 | 0.8 | 0.424 |

**Table S29: Summary of the selected model for CHG 13 for worldwide data.** φ correspond to the estimated time autoregression coefficient. ν and ρ are respectively the estimated smoothness and scaling parameters of the Matérn correlation function. *: p<0.05, **: p<0.01, ***: p<0.001.

$\varphi = 0.914$, $\nu = 6.33 \times 10^{-3}$, $\rho = 0.303$

| Variable | Estimate | Conditional standard error | t | p |
|---|---|---|---|---|
| Intercept | -24.5 | $3.48 \times 10^{3}$ | $-7.03 \times 10^{-3}$ | 0.994 |
| Domestic animals | $-2.15 \times 10^{2}$ | $2.63 \times 10^{6}$ | $-8.16 \times 10^{-5}$ | 1 |
| Farms | 17.3 | $3.48 \times 10^{3}$ | $4.97 \times 10^{-3}$ | 0.996 |
| Flora, fauna | 17.1 | $3.48 \times 10^{3}$ | $4.91 \times 10^{-3}$ | 0.996 |
| Human samples | 18.4 | $3.48 \times 10^{3}$ | $5.29 \times 10^{-3}$ | 0.996 |
| Human habitat | 3.44 | $3.48 \times 10^{3}$ | $9.88 \times 10^{-4}$ | 0.999 |
| Freshwater | $-2.74 \times 10^{2}$ | $2.43 \times 10^{6}$ | $-1.13 \times 10^{-4}$ | 1 |
| Clinical samples | 18.6 | $3.48 \times 10^{3}$ | $5.34 \times 10^{-3}$ | 0.996 |
| Sludge, waste | 19.9 | $3.48 \times 10^{3}$ | $5.73 \times 10^{-3}$ | 0.995 |
| Soil | 7.25 | $3.48 \times 10^{3}$ | $2.08 \times 10^{-3}$ | 0.998 |
| Trade | -0.847 | $3.85 \times 10^{3}$ | $-2.2 \times 10^{-4}$ | 1 |
| Migration | -0.154 | $2.48 \times 10^{3}$ | $-6.2 \times 10^{-5}$ | 1 |
| Domestic animals × Trade | $-3.48 \times 10^{2}$ | $2.26 \times 10^{6}$ | $-1.54 \times 10^{-4}$ | 1 |
| Farms × Trade | -0.535 | $3.85 \times 10^{3}$ | $-1.39 \times 10^{-4}$ | 1 |
| Flora, fauna × Trade | -1.36 | $3.85 \times 10^{3}$ | $-3.54 \times 10^{-4}$ | 1 |
| Human samples × Trade | 0.598 | $3.85 \times 10^{3}$ | $1.55 \times 10^{-4}$ | 1 |
| Human habitat × Trade | 0.8 | $3.85 \times 10^{3}$ | $2.08 \times 10^{-4}$ | 1 |
| Freshwater × Trade | 4.81 | $2.59 \times 10^{6}$ | $1.86 \times 10^{-6}$ | 1 |
| Clinical samples × Trade | $9.19 \times 10^{-2}$ | $3.85 \times 10^{3}$ | $2.39 \times 10^{-5}$ | 1 |
| Sludge, waste × Trade | -0.213 | $3.85 \times 10^{3}$ | $-5.53 \times 10^{-5}$ | 1 |
| Soil × Trade | 1.3 | $3.85 \times 10^{3}$ | $3.37 \times 10^{-4}$ | 1 |
| Domestic animals × Migration | $1.36 \times 10^{2}$ | $1.69 \times 10^{6}$ | $8.07 \times 10^{-5}$ | 1 |
| Farms × Migration | -3.3 | $2.48 \times 10^{3}$ | $-1.33 \times 10^{-3}$ | 0.999 |
| Flora, fauna × Migration | -3.74 | $2.48 \times 10^{3}$ | $-1.51 \times 10^{-3}$ | 0.999 |
| Human samples × Migration | 0.108 | $2.48 \times 10^{3}$ | $4.36 \times 10^{-5}$ | 1 |
| Human habitat × Migration | -48.6 | $2.48 \times 10^{3}$ | $-1.96 \times 10^{-2}$ | 0.984 |
| Freshwater × Migration | 1.16 | $2.65 \times 10^{6}$ | $4.39 \times 10^{-7}$ | 1 |
| Clinical samples × Migration | 0.192 | $2.48 \times 10^{3}$ | $7.71 \times 10^{-5}$ | 1 |
| Sludge, waste × Migration | -0.161 | $2.48 \times 10^{3}$ | $-6.47 \times 10^{-5}$ | 1 |
| Soil × Migration | -31.6 | $2.48 \times 10^{3}$ | $-1.27 \times 10^{-2}$ | 0.99 |

**Table S30: Summary of the selected model for CHG 14 for worldwide data.** φ correspond to the estimated time autoregression coefficient. ν and ρ are respectively the estimated smoothness and scaling parameters of the Matérn correlation function. *: p<0.05, **: p<0.01, ***: p<0.001.

φ = 0.294, ν = 16.7, ρ = 0.922

| **Variable** | **Estimate** | **Conditional standard error** | **t** | **p** |
|---|---|---|---|---|
| Intercept | -8.94 | 1.92 | -4.66 | $3.23 \times 10^{-6}$ *** |
| Domestic animals | -29 | $4.39 \times 10^{6}$ | $-6.6 \times 10^{-6}$ | 1 |
| Farms | 2.93 | 1.87 | 1.57 | 0.117 |
| Flora, fauna | 0.685 | 3.3 | 0.207 | 0.836 |
| Human samples | -5.35 | 3.29 | -1.63 | 0.104 |
| Human habitat | -28.4 | $3.88 \times 10^{6}$ | $-7.33 \times 10^{-6}$ | 1 |
| Freshwater | -28.9 | $2.55 \times 10^{6}$ | $-1.13 \times 10^{-5}$ | 1 |
| Clinical samples | 1.11 | 1.87 | 0.591 | 0.555 |
| Sludge, waste | -29.2 | $5.48 \times 10^{6}$ | $-5.32 \times 10^{-6}$ | 1 |
| Soil | -28.4 | $2.13 \times 10^{6}$ | $-1.34 \times 10^{-5}$ | 1 |
| Trade | -0.984 | 1.55 | -0.635 | 0.526 |
| Migration | -1.73 | 6.3 | -0.275 | 0.783 |
| Domestic animals × Trade | 0.468 | $2.76 \times 10^{6}$ | $1.69 \times 10^{-7}$ | 1 |
| Farms × Trade | 0.83 | 1.55 | 0.537 | 0.591 |
| Flora, fauna × Trade | -0.228 | 3.42 | $-6.67 \times 10^{-2}$ | 0.947 |
| Human samples × Trade | 1.27 | 1.57 | 0.806 | 0.42 |
| Human habitat × Trade | 0.426 | $2.01 \times 10^{6}$ | $2.12 \times 10^{-7}$ | 1 |
| Freshwater × Trade | 0.504 | $1.44 \times 10^{6}$ | $3.5 \times 10^{-7}$ | 1 |
| Clinical samples × Trade | 0.269 | 1.56 | 0.173 | 0.862 |
| Sludge, waste × Trade | 0.452 | $2.4 \times 10^{6}$ | $1.88 \times 10^{-7}$ | 1 |
| Soil × Trade | 0.197 | $2 \times 10^{6}$ | $9.86 \times 10^{-8}$ | 1 |
| Domestic animals × Migration | 1.63 | $2.67 \times 10^{6}$ | $6.11 \times 10^{-7}$ | 1 |
| Farms × Migration | 1.76 | 6.3 | 0.28 | 0.78 |
| Flora, fauna × Migration | -0.181 | 12 | $-1.5 \times 10^{-2}$ | 0.988 |
| Human samples × Migration | -21 | 10.3 | -2.05 | $4.08 \times 10^{-2}$ * |
| Human habitat × Migration | 1.38 | $5.88 \times 10^{6}$ | $2.34 \times 10^{-7}$ | 1 |
| Freshwater × Migration | 1.36 | $3.21 \times 10^{6}$ | $4.25 \times 10^{-7}$ | 1 |
| Clinical samples × Migration | 1.4 | 6.3 | 0.223 | 0.824 |
| Sludge, waste × Migration | 1.65 | $4.24 \times 10^{6}$ | $3.9 \times 10^{-7}$ | 1 |
| Soil × Migration | 1.62 | $3.35 \times 10^{6}$ | $4.83 \times 10^{-7}$ | 1 |

**Table S31: Summary of the selected model for CHG 15 for worldwide data.** φ correspond to the estimated time autoregression coefficient. ν and ρ are respectively the estimated smoothness and scaling parameters of the Matérn correlation function. *: p<0.05, **: p<0.01, ***: p<0.001.

$\varphi = 2.17 \times 10^{-2}$, $\nu = 16.7$, $\rho = 0.724$

| **Variable** | **Estimate** | **Conditional standard error** | **t** | **p** |
|---|---|---|---|---|
| Intercept | -5.96 | 0.603 | -9.88 | 0 *** |
| Domestic animals | -31.3 | $3.91 \times 10^{6}$ | $-8.01 \times 10^{-6}$ | 1 |
| Farms | -1.63 | 0.648 | -2.51 | $1.2 \times 10^{-2}$ * |
| Flora, fauna | 0.809 | 0.701 | 1.15 | 0.248 |
| Human samples | -31.3 | $8.32 \times 10^{5}$ | $-3.76 \times 10^{-5}$ | 1 |
| Human habitat | $-9.06 \times 10^{-2}$ | 0.902 | -0.1 | 0.92 |
| Freshwater | -31.2 | $2.34 \times 10^{6}$ | $-1.33 \times 10^{-5}$ | 1 |
| Clinical samples | -4.82 | 1.15 | -4.19 | $2.74 \times 10^{-5}$ *** |
| Sludge, waste | -31.1 | $3.57 \times 10^{6}$ | $-8.7 \times 10^{-6}$ | 1 |
| Soil | 1.26 | 0.528 | 2.39 | $1.71 \times 10^{-2}$ * |
| Trade | -0.221 | 0.229 | -0.968 | 0.333 |
| Migration | -0.24 | 0.18 | -1.34 | 0.181 |

**Table S32: Summary of the selected model for CHG 17.1 for worldwide data.** φ correspond to the estimated time autoregression coefficient. ν and ρ are respectively the estimated smoothness and scaling parameters of the Matérn correlation function. *: $p<0.05$, **: $p<0.01$, ***: $p<0.001$.

$\varphi = 0.704$, $\nu = 5.7 \times 10^{-3}$, $\rho = 0.178$

| **Variable** | **Estimate** | **Conditional standard error** | **t** | **p** |
|---|---|---|---|---|
| Intercept | -36.9 | $3.13 \times 10^{6}$ | $-1.18 \times 10^{-5}$ | 1 |
| Domestic animals | -7.7 | $5 \times 10^{6}$ | $-1.54 \times 10^{-6}$ | 1 |
| Farms | -7.01 | $3.25 \times 10^{6}$ | $-2.16 \times 10^{-6}$ | 1 |
| Flora, fauna | -7.1 | $4.95 \times 10^{6}$ | $-1.43 \times 10^{-6}$ | 1 |
| Human samples | 25.1 | $3.13 \times 10^{6}$ | $8.03 \times 10^{-6}$ | 1 |
| Human habitat | 29.3 | $3.13 \times 10^{6}$ | $9.37 \times 10^{-6}$ | 1 |
| Freshwater | 27.9 | $3.13 \times 10^{6}$ | $8.93 \times 10^{-6}$ | 1 |
| Clinical samples | 27.7 | $3.13 \times 10^{6}$ | $8.87 \times 10^{-6}$ | 1 |
| Sludge, waste | 27.7 | $3.13 \times 10^{6}$ | $8.87 \times 10^{-6}$ | 1 |
| Soil | -6.69 | $3.68 \times 10^{6}$ | $-1.82 \times 10^{-6}$ | 1 |
| Trade | 0.163 | 0.134 | 1.22 | 0.224 |
| Migration | -2.89 | 0.862 | -3.35 | $8.09 \times 10^{-4}$ *** |

**Table S33: Summary of the selected model for CHG 19 for worldwide data.** φ correspond to the estimated time autoregression coefficient. ν and ρ are respectively the estimated smoothness and scaling parameters of the Matérn correlation function. *: p<0.05, **: p<0.01, ***: p<0.001.

$\varphi = 0.931$, $\nu = 1.88 \times 10^{-2}$, $\rho = 1.89$

| Variable | Estimate | Conditional standard error | t | p |
|---|---|---|---|---|
| Intercept | -15.4 | 10.6 | -1.45 | 0.148 |
| Domestic animals | 8.65 | 10.6 | 0.816 | 0.414 |
| Farms | 9.64 | 10.6 | 0.912 | 0.362 |
| Flora, fauna | 10.6 | 10.6 | 1 | 0.316 |
| Human samples | 9.07 | 10.6 | 0.858 | 0.391 |
| Human habitat | 7.88 | 10.6 | 0.741 | 0.458 |
| Freshwater | -23.1 | $2.68 \times 10^{6}$ | $-8.61 \times 10^{-6}$ | 1 |
| Clinical samples | 10.2 | 10.6 | 0.965 | 0.335 |
| Sludge, waste | 10.5 | 10.6 | 0.989 | 0.323 |
| Soil | 0.975 | 12.8 | $7.62 \times 10^{-2}$ | 0.939 |
| Trade | 1.21 | 2.44 | 0.495 | 0.621 |
| Migration | 0.582 | 0.364 | 1.6 | 0.11 |
| Domestic animals × Trade | -0.813 | 2.47 | -0.329 | 0.742 |
| Farms × Trade | -1.61 | 2.45 | -0.66 | 0.509 |
| Flora, fauna × Trade | -1.68 | 2.47 | -0.679 | 0.497 |
| Human samples × Trade | -1.27 | 2.44 | -0.519 | 0.604 |
| Human habitat × Trade | -0.933 | 2.52 | -0.371 | 0.711 |
| Freshwater × Trade | -1.93 | $2.74 \times 10^{6}$ | $-7.06 \times 10^{-7}$ | 1 |
| Clinical samples × Trade | -1.42 | 2.44 | -0.58 | 0.562 |
| Sludge, waste × Trade | -1.26 | 2.45 | -0.514 | 0.607 |
| Soil × Trade | -4 | 4.5 | -0.889 | 0.374 |
| Domestic animals × Migration | -0.717 | 0.417 | -1.72 | $8.57 \times 10^{-2}$ * |
| Farms × Migration | -1.27 | 0.409 | -3.11 | $1.86 \times 10^{-3}$ ** |
| Flora, fauna × Migration | -1.58 | 0.614 | -2.57 | $1.02 \times 10^{-2}$ * |
| Human samples × Migration | -0.499 | 0.374 | -1.34 | 0.182 |
| Human habitat × Migration | -0.496 | 0.601 | -0.826 | 0.409 |
| Freshwater × Migration | -0.385 | $2.94 \times 10^{6}$ | $-1.31 \times 10^{-7}$ | 1 |
| Clinical samples × Migration | -0.702 | 0.365 | -1.93 | $5.41 \times 10^{-2}$ * |
| Sludge, waste × Migration | -0.682 | 0.411 | -1.66 | $9.68 \times 10^{-2}$ * |
| Soil × Migration | -12.4 | 12.9 | -0.96 | 0.337 |

**Table S34: Summary of the selected model for CHG 20 for worldwide data.** φ correspond to the estimated time autoregression coefficient. ν and ρ are respectively the estimated smoothness and scaling parameters of the Matérn correlation function. *: p<0.05, **: p<0.01, ***: p<0.001.

φ = 0.283, ν = 16.7, ρ = 0.823

| **Variable** | **Estimate** | **Conditional standard error** | **t** | **p** |
|---|---|---|---|---|
| Intercept | -11 | 2.33 | -4.69 | $2.67 \times 10^{-6}$ *** |
| Domestic animals | -26.9 | $4.38 \times 10^{6}$ | $-6.14 \times 10^{-6}$ | 1 |
| Farms | 4.94 | 2.27 | 2.17 | $2.96 \times 10^{-2}$ * |
| Flora, fauna | 2.49 | 4.18 | 0.595 | 0.552 |
| Human samples | $7.31 \times 10^{-2}$ | 2.67 | $2.74 \times 10^{-2}$ | 0.978 |
| Human habitat | -26.4 | $3.88 \times 10^{6}$ | $-6.8 \times 10^{-6}$ | 1 |
| Freshwater | -26.9 | $2.53 \times 10^{6}$ | $-1.06 \times 10^{-5}$ | 1 |
| Clinical samples | 3.34 | 2.28 | 1.46 | 0.144 |
| Sludge, waste | -27.1 | $5.52 \times 10^{6}$ | $-4.9 \times 10^{-6}$ | 1 |
| Soil | -26.6 | $2.1 \times 10^{6}$ | $-1.27 \times 10^{-5}$ | 1 |
| Trade | 0.289 | 0.493 | 0.586 | 0.558 |
| Migration | -10.8 | 7.07 | -1.52 | 0.128 |
| Domestic animals × Trade | -0.844 | $2.89 \times 10^{6}$ | $-2.92 \times 10^{-7}$ | 1 |
| Farms × Trade | -0.413 | 0.492 | -0.84 | 0.401 |
| Flora, fauna × Trade | -1.65 | 3.5 | -0.471 | 0.638 |
| Human samples × Trade | -0.289 | 0.533 | -0.543 | 0.587 |
| Human habitat × Trade | -0.84 | $2.03 \times 10^{6}$ | $-4.14 \times 10^{-7}$ | 1 |
| Freshwater × Trade | -0.772 | $1.45 \times 10^{6}$ | $-5.33 \times 10^{-7}$ | 1 |
| Clinical samples × Trade | -1.03 | 0.531 | -1.93 | $5.31 \times 10^{-2}$ * |
| Sludge, waste × Trade | -0.85 | $2.45 \times 10^{6}$ | $-3.47 \times 10^{-7}$ | 1 |
| Soil × Trade | -1.06 | $2.08 \times 10^{6}$ | $-5.1 \times 10^{-7}$ | 1 |
| Domestic animals × Migration | 10.6 | $2.74 \times 10^{6}$ | $3.86 \times 10^{-6}$ | 1 |
| Farms × Migration | 10.8 | 7.07 | 1.53 | 0.127 |
| Flora, fauna × Migration | 7.83 | 14.1 | 0.556 | 0.578 |
| Human samples × Migration | -1.99 | 8.09 | -0.246 | 0.806 |
| Human habitat × Migration | 10.2 | $5.3 \times 10^{6}$ | $1.92 \times 10^{-6}$ | 1 |
| Freshwater × Migration | 10.2 | $3.1 \times 10^{6}$ | $3.28 \times 10^{-6}$ | 1 |
| Clinical samples × Migration | 10.2 | 7.07 | 1.45 | 0.148 |
| Sludge, waste × Migration | 10.5 | $4.13 \times 10^{6}$ | $2.55 \times 10^{-6}$ | 1 |
| Soil × Migration | 10.5 | $3.27 \times 10^{6}$ | $3.2 \times 10^{-6}$ | 1 |

**Table S35: Summary of the selected model for CHG 23 for worldwide data.** φ correspond to the estimated time autoregression coefficient. ν and ρ are respectively the estimated smoothness and scaling parameters of the Matérn correlation function. *: $p<0.05$, **: $p<0.01$, ***: $p<0.001$.

φ = 0.521, ν = 1.22, ρ = 0.534

| Variable | Estimate | Conditional standard error | t | p |
|---|---|---|---|---|
| Intercept | -25.7 | $4.1 \times 10^{3}$ | $-6.27 \times 10^{-3}$ | 0.995 |
| Domestic animals | $-4.01 \times 10^{2}$ | $4.8 \times 10^{6}$ | $-8.36 \times 10^{-5}$ | 1 |
| Farms | $-4.01 \times 10^{2}$ | $1.09 \times 10^{6}$ | $-3.66 \times 10^{-4}$ | 1 |
| Flora, fauna | $-4 \times 10^{2}$ | $4.13 \times 10^{6}$ | $-9.69 \times 10^{-5}$ | 1 |
| Human samples | 18.6 | $4.1 \times 10^{3}$ | $4.54 \times 10^{-3}$ | 0.996 |
| Human habitat | -29.7 | $4.1 \times 10^{3}$ | $-7.23 \times 10^{-3}$ | 0.994 |
| Freshwater | 17 | $4.1 \times 10^{3}$ | $4.13 \times 10^{-3}$ | 0.997 |
| Clinical samples | 19.8 | $4.1 \times 10^{3}$ | $4.84 \times 10^{-3}$ | 0.996 |
| Sludge, waste | $-2.03 \times 10^{2}$ | $6.25 \times 10^{4}$ | $-3.24 \times 10^{-3}$ | 0.997 |
| Soil | 12.9 | $4.1 \times 10^{3}$ | $3.13 \times 10^{-3}$ | 0.997 |
| Trade | -0.717 | $3.22 \times 10^{3}$ | $-2.23 \times 10^{-4}$ | 1 |
| Migration | 0.102 | $2.9 \times 10^{3}$ | $3.53 \times 10^{-5}$ | 1 |
| Domestic animals × Trade | 4.19 | $3.26 \times 10^{6}$ | $1.29 \times 10^{-6}$ | 1 |
| Farms × Trade | 2.46 | $5.65 \times 10^{5}$ | $4.35 \times 10^{-6}$ | 1 |
| Flora, fauna × Trade | 1.5 | $2.35 \times 10^{6}$ | $6.38 \times 10^{-7}$ | 1 |
| Human samples × Trade | 0.91 | $3.22 \times 10^{3}$ | $2.83 \times 10^{-4}$ | 1 |
| Human habitat × Trade | 2.83 | $3.22 \times 10^{3}$ | $8.79 \times 10^{-4}$ | 0.999 |
| Freshwater × Trade | -0.351 | $3.22 \times 10^{3}$ | $-1.09 \times 10^{-4}$ | 1 |
| Clinical samples × Trade | 0.423 | $3.22 \times 10^{3}$ | $1.31 \times 10^{-4}$ | 1 |
| Sludge, waste × Trade | $-2.58 \times 10^{2}$ | $1.57 \times 10^{5}$ | $-1.64 \times 10^{-3}$ | 0.999 |
| Soil × Trade | -6.99 | $3.22 \times 10^{3}$ | $-2.17 \times 10^{-3}$ | 0.998 |
| Domestic animals × Migration | -0.821 | $2.08 \times 10^{6}$ | $-3.95 \times 10^{-7}$ | 1 |
| Farms × Migration | -0.64 | $3.4 \times 10^{5}$ | $-1.88 \times 10^{-6}$ | 1 |
| Flora, fauna × Migration | $6.75 \times 10^{-2}$ | $2.53 \times 10^{6}$ | $2.67 \times 10^{-8}$ | 1 |
| Human samples × Migration | -2.87 | $2.9 \times 10^{3}$ | $-9.9 \times 10^{-4}$ | 0.999 |
| Human habitat × Migration | $-1.63 \times 10^{2}$ | $2.91 \times 10^{3}$ | $-5.62 \times 10^{-2}$ | 0.955 |
| Freshwater × Migration | -1.54 | $2.9 \times 10^{3}$ | $-5.29 \times 10^{-4}$ | 1 |
| Clinical samples × Migration | -0.195 | $2.9 \times 10^{3}$ | $-6.71 \times 10^{-5}$ | 1 |
| Sludge, waste × Migration | $-1.94 \times 10^{2}$ | $3.11 \times 10^{5}$ | $-6.24 \times 10^{-4}$ | 1 |
| Soil × Migration | -5.92 | $2.9 \times 10^{3}$ | $-2.04 \times 10^{-3}$ | 0.998 |

**Table S36: Summary of the selected model for CHG 25 for worldwide data.** φ correspond to the estimated time autoregression coefficient. ν and ρ are respectively the estimated smoothness and scaling parameters of the Matérn correlation function. *: $p<0.05$, **: $p<0.01$, ***: $p<0.001$.

$\varphi = -0.469$, $\nu = 5.01 \times 10^{-3}$, $\rho = 2.54 \times 10^{-2}$

| Variable | Estimate | Conditional standard error | t | p |
|---|---|---|---|---|
| Intercept | -39.6 | 25.7 | -1.54 | 0.123 |
| Trade | -28.9 | 36.1 | -0.801 | 0.423 |
| Migration | $-1.23 \times 10^{2}$ | $1.43 \times 10^{2}$ | -0.856 | 0.392 |

**Table S37: Summary of the selected model for CHG 27 for worldwide data.** φ correspond to the estimated time autoregression coefficient. ν and ρ are respectively the estimated smoothness and scaling parameters of the Matérn correlation function. *: p<0.05, **: p<0.01, ***: p<0.001.

$\varphi = 1.14 \times 10^{-2}$, $\nu = 8.21 \times 10^{-2}$, $\rho = 1.29 \times 10^{-2}$

| **Variable** | **Estimate** | **Conditional standard error** | **t** | **p** |
|---|---|---|---|---|
| Intercept | -4.61 | 0.549 | -8.39 | 0 *** |
| Domestic animals | -32.5 | $4.54 \times 10^{6}$ | $-7.15 \times 10^{-6}$ | 1 |
| Farms | -2.1 | 0.509 | -4.12 | $3.76 \times 10^{-5}$ *** |
| Flora, fauna | -0.698 | 0.85 | -0.821 | 0.412 |
| Human samples | -1.45 | 0.317 | -4.58 | $4.66 \times 10^{-6}$ *** |
| Human habitat | -3.63 | 3.63 | -0.999 | 0.318 |
| Freshwater | -0.198 | 0.367 | -0.541 | 0.589 |
| Clinical samples | -0.395 | 0.209 | -1.89 | $5.84 \times 10^{-2}$ * |
| Sludge, waste | -0.156 | 0.659 | -0.236 | 0.813 |
| Soil | -0.437 | 0.389 | -1.12 | 0.261 |
| Trade | $-5.93 \times 10^{-2}$ | $8.65 \times 10^{-2}$ | -0.686 | 0.493 |
| Migration | 0.651 | $8.8 \times 10^{-2}$ | 7.39 | $1.45 \times 10^{-13}$ *** |
| Domestic animals × Trade | -0.464 | $3.5 \times 10^{6}$ | $-1.33 \times 10^{-7}$ | 1 |
| Farms × Trade | $6.4 \times 10^{-2}$ | 0.119 | 0.539 | 0.59 |
| Flora, fauna × Trade | -0.198 | 0.447 | -0.444 | 0.657 |
| Human samples × Trade | -0.483 | 0.268 | -1.8 | $7.16 \times 10^{-2}$ * |
| Human habitat × Trade | $2.75 \times 10^{-2}$ | 0.319 | $8.63 \times 10^{-2}$ | 0.931 |
| Freshwater × Trade | -1.38 | 0.716 | -1.92 | $5.47 \times 10^{-2}$ * |
| Clinical samples × Trade | -1.09 | 0.175 | -6.26 | $3.79 \times 10^{-10}$ *** |
| Sludge, waste × Trade | -0.256 | 0.284 | -0.901 | 0.368 |
| Soil × Trade | -0.323 | 0.322 | -1 | 0.316 |
| Domestic animals × Migration | -0.421 | $3.61 \times 10^{6}$ | $-1.17 \times 10^{-7}$ | 1 |
| Farms × Migration | -2.99 | 1.08 | -2.76 | $5.82 \times 10^{-3}$ ** |
| Flora, fauna × Migration | -0.776 | 0.7 | -1.11 | 0.268 |
| Human samples × Migration | -1.02 | 0.312 | -3.27 | $1.08 \times 10^{-3}$ ** |
| Human habitat × Migration | -10.2 | 8.7 | -1.17 | 0.243 |
| Freshwater × Migration | -1.05 | 0.426 | -2.47 | $1.33 \times 10^{-2}$ * |
| Clinical samples × Migration | -0.883 | 0.148 | -5.95 | $2.65 \times 10^{-9}$ *** |
| Sludge, waste × Migration | -0.665 | 0.477 | -1.4 | 0.163 |
| Soil × Migration | $-7.66 \times 10^{-3}$ | 0.306 | $-2.51 \times 10^{-2}$ | 0.98 |

**Table S39: Impact of the association of AMEGs with mobile genetic elements with their duplication risk**

| Minimum identity between genomic contexts | Minimum identity between genes | Odds ratio | 500-permutation p-value |
|---|---|---|---|
| 50% | 80% | 14.2 | 0.001 |
| 50% | 85% | 11.9 | 0.001 |
| 50% | 90% | 10.7 | 0.001 |
| 50% | 95% | 10.4 | 0.001 |
| 70% | 80% | 14.2 | 0.001 |
| 70% | 85% | 11.9 | 0.001 |
| 70% | 90% | 10.7 | 0.001 |
| 70% | 95% | 10.4 | 0.001 |
| 80% | 80% | 14.2 | 0.001 |
| 0.001 | 85% | 11.9 | 0.001 |
| 0.001 | 90% | 10.7 | 0.001 |
| 80% | 95% | 10.4 | 0.001 |
| 90% | 80% | 14.2 | 0.001 |
| 90% | 85% | 11.9 | 0.001 |
| 90% | 90% | 10.7 | 0.001 |
| 90% | 95% | 10.4 | 0.001 |

**Table S40: Impact of the association of AMEGs with different types of mobile genetic elements with their duplication risk**

| Minimum identity between genomic contexts | Minimum identity between genes | Type of mobile genetic element | Odds ratio | 500-permutation p-value |
|---|---|---|---|---|
| 50% | 80% | Intergenomic mobility | 18.5 | 0.001 |
| 50% | 85% | Intergenomic mobility | 14.7 | 0.001 |
| 50% | 90% | Intergenomic mobility | 13.7 | 0.001 |
| 50% | 95% | Intergenomic mobility | 13.3 | 0.001 |
| 70% | 80% | Intergenomic mobility | 18.5 | 0.001 |
| 70% | 85% | Intergenomic mobility | 14.7 | 0.001 |
| 70% | 90% | Intergenomic mobility | 13.7 | 0.001 |
| 70% | 95% | Intergenomic mobility | 13.3 | 0.001 |
| 80% | 80% | Intergenomic mobility | 18.5 | 0.001 |
| 80% | 85% | Intergenomic mobility | 14.7 | 0.001 |
| 80% | 90% | Intergenomic mobility | 13.7 | 0.001 |
| 80% | 95% | Intergenomic mobility | 13.3 | 0.001 |
| 90% | 80% | Intergenomic mobility | 18.5 | 0.001 |
| 90% | 85% | Intergenomic mobility | 14.7 | 0.001 |
| 90% | 90% | Intergenomic mobility | 13.7 | 0.001 |
| 90% | 95% | Intergenomic mobility | 13.3 | 0.001 |
| 50% | 80% | Intragenomic mobility | 12.0 | 0.001 |
| 50% | 85% | Intragenomic mobility | 12.0 | 0.002 |
| 50% | 90% | Intragenomic mobility | 12.0 | 0.002 |
| 50% | 95% | Intragenomic mobility | 10.8 | 0.003 |
| 70% | 80% | Intragenomic mobility | 12.0 | 0.001 |
| 70% | 85% | Intragenomic mobility | 12.0 | 0.002 |
| 70% | 90% | Intragenomic mobility | 12.0 | 0.002 |
| 70% | 95% | Intragenomic mobility | 10.8 | 0.003 |
| 80% | 80% | Intragenomic mobility | 12.0 | 0.001 |
| 80% | 85% | Intragenomic mobility | 12.0 | 0.002 |
| 80% | 90% | Intragenomic mobility | 12.0 | 0.002 |
| 80% | 95% | Intragenomic mobility | 10.8 | 0.003 |
| 90% | 80% | Intragenomic mobility | 12.0 | 0.001 |
| 90% | 85% | Intragenomic mobility | 12.0 | 0.002 |
| 90% | 90% | Intragenomic mobility | 12.0 | 0.002 |
| 90% | 95% | Intragenomic mobility | 10.8 | 0.003 |
| 50% | 80% | Intra- and intergenomic mobility | 16.0 | 0.001 |
| 50% | 85% | Intra- and intergenomic mobility | 14.1 | 0.001 |
| 50% | 90% | Intra- and intergenomic mobility | 12.1 | 0.001 |
| 50% | 95% | Intra- and intergenomic mobility | 12.1 | 0.001 |
| 70% | 80% | Intra- and intergenomic mobility | 16.0 | 0.001 |
| 70% | 85% | Intra- and intergenomic mobility | 14.1 | 0.001 |
| 70% | 90% | Intra- and intergenomic mobility | 12.1 | 0.001 |
| 70% | 95% | Intra- and intergenomic mobility | 12.1 | 0.001 |
| 80% | 80% | Intra- and intergenomic mobility | 16.0 | 0.001 |
| 80% | 85% | Intra- and intergenomic mobility | 14.1 | 0.001 |
| 80% | 90% | Intra- and intergenomic mobility | 12.1 | 0.001 |
| 80% | 95% | Intra- and intergenomic mobility | 12.1 | 0.001 |
| 90% | 80% | Intra- and intergenomic mobility | 16.0 | 0.001 |
| 90% | 85% | Intra- and intergenomic mobility | 14.1 | 0.001 |
| 90% | 90% | Intra- and intergenomic mobility | 12.1 | 0.001 |
| 90% | 95% | Intra- and intergenomic mobility | 12.1 | 0.001 |

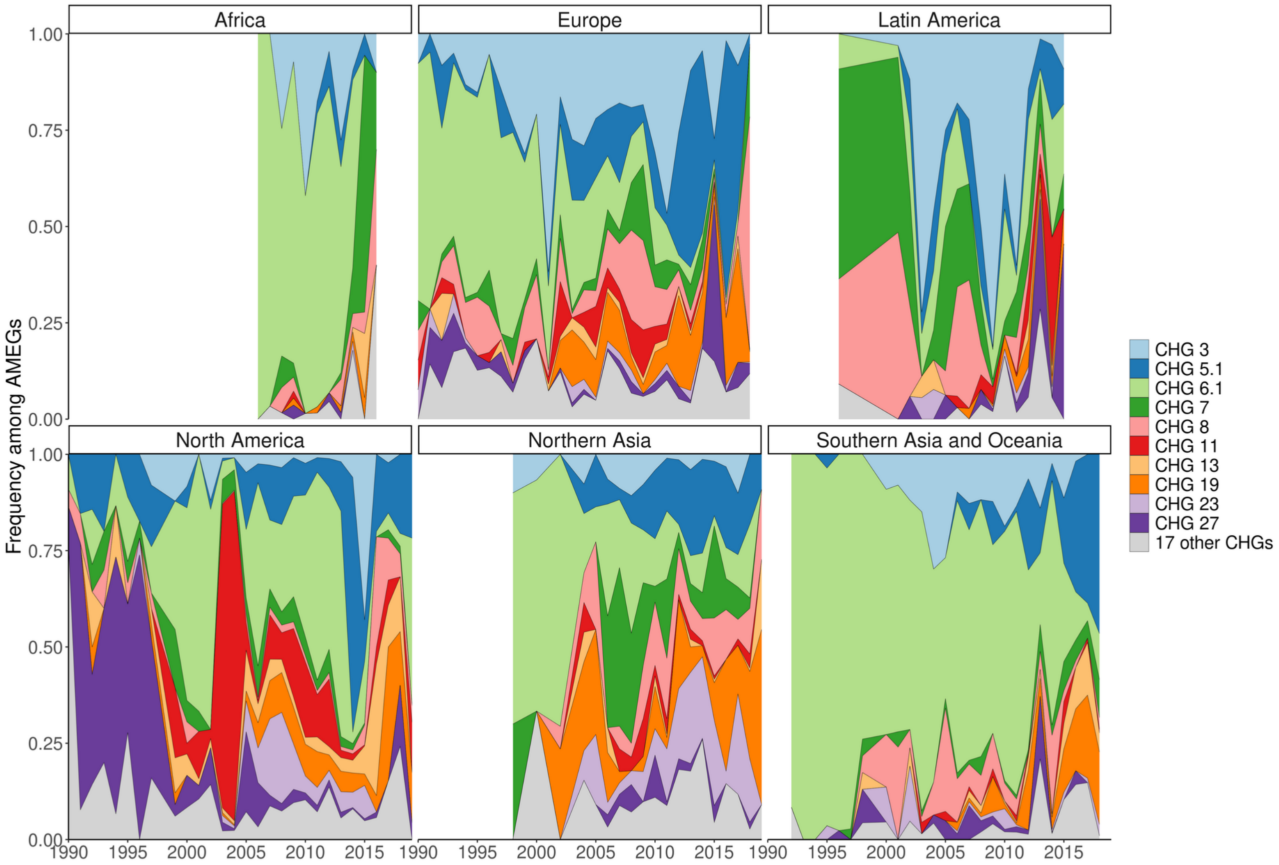


**Figure S1: Evolution of continental CHG frequencies among sampled AMEGs between 1990 and 2019.** IMAGE24 regions were gathered to higher continental ensembles in order to more representative frequencies. Western Europe, Central Europe, and “Ukraine+” were gathered to Europe; Canada and the USA to North America; the rest of Americas to Latin America; all African landmasses to Africa; Oceania, Southeastern Asia, “Indonesia+”, and “India+” to Southern Asia and Oceania; and Korea, Japan, Middle East, Turkey, “China+”, “Asia-Stan”, and “Russia+” to Northern Asia. CHG frequencies were only computed when at least 10 ACBs had been sampled in a continent in a year.

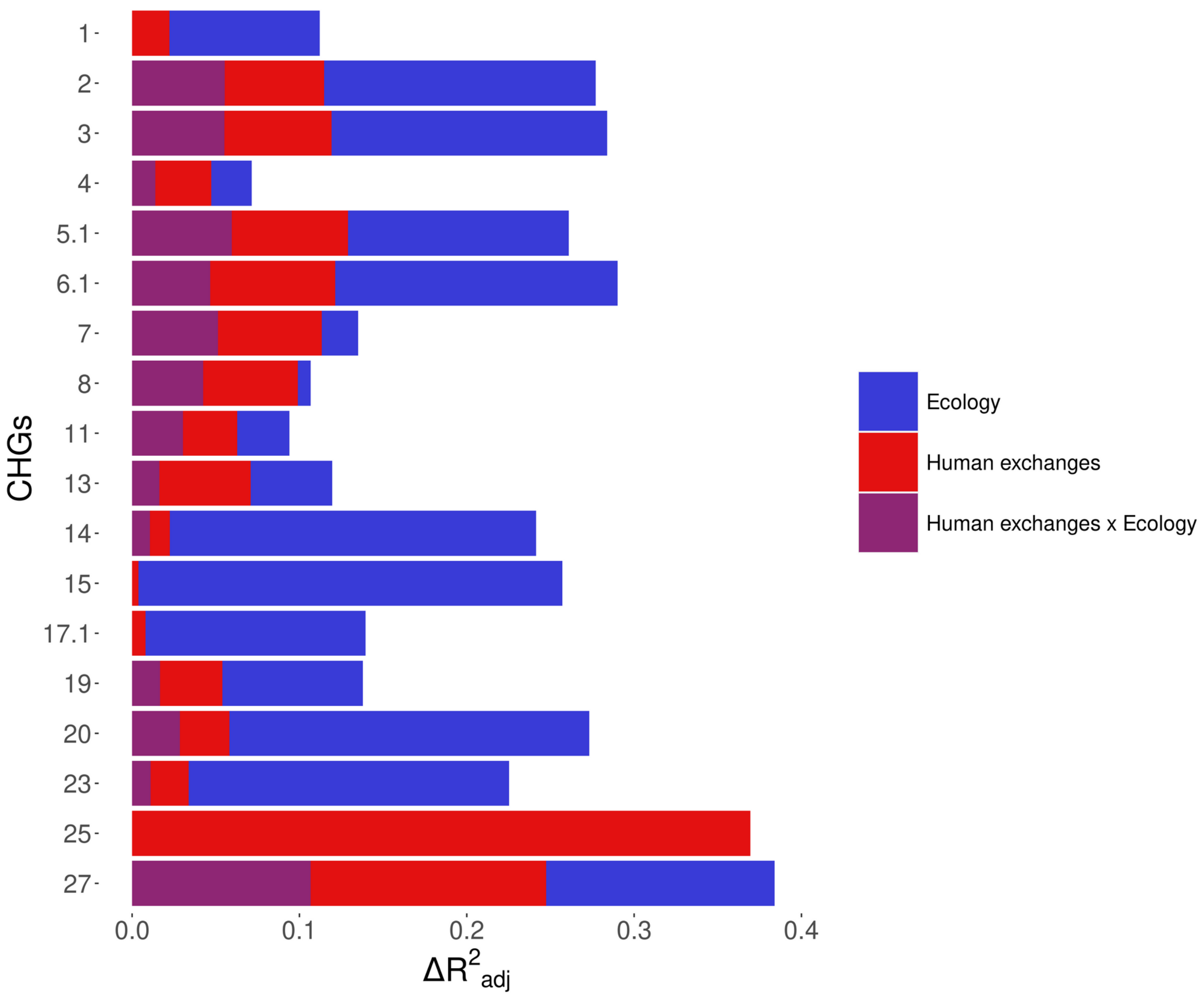


**Figure S2: Relative importance of several factors to explain the distribution of aminoglycoside-resistant bacteria in the world between 1997 and 2018.** Logistic regressions with a spatial Matérn correlation structure were computed to explain the frequency of 18 CHGs in samples. This figure represents the contribution of each variable in each selected model, as the fraction of adjusted McFadden's pseudo-R2 explained by adding this variable.
